# Morning Glucagon Disrupts Insulin Induced Hepatic Metabolic Memory and Subsequent Afternoon Glucose Metabolism in Canines

**DOI:** 10.1101/2025.02.25.639957

**Authors:** Hannah L. Waterman, Marta S. Smith, Ben Farmer, Kalisha Yankey, Karin J. Bosma, Richard M. O’Brien, Derek P. Claxton, Tristan Howard, Guillaume Kraft, Alan D. Cherrington, Dale S. Edgerton

**Author notes:** **Corresponding author information: Name:** Hannah Lynn Waterman, PhD **Current Affiliation:** Duke University **Email:**.

## Abstract

The Staub-Traugott effect, or second-meal phenomenon, describes improved glucose disposal after a second identical meal. We previously showed that morning hyperinsulinemia primes the liver to enhance afternoon net hepatic glucose uptake and glycogen storage. However, mixed meals trigger co-secretion of insulin and glucagon, and glucagon is traditionally viewed as opposing insulin’s hepatic actions. Whether glucagon modifies the persistence of insulin’s priming effects across sequential metabolic challenges is unknown. Therefore, we investigated whether morning hyperglucagonemia alters the ability of morning hyperinsulinemia to prime subsequent hepatic glucose metabolism. Conscious dogs underwent two pancreatic clamp periods separated by a 1.5h rest period. Endogenous insulin and glucagon were suppressed with somatostatin and replaced intraportally at defined rates. During a 4h morning hyperinsulinemic-euglycemic clamp, dogs received matched insulin prime infusions with either basal glucagon (AM INS; n=8) or elevated glucagon (AM INS+GCG; n=8). After the rest period, both groups underwent a 2.5h afternoon hyperinsulinemic-hyperglycemic clamp under identical hormonal conditions. Afternoon net hepatic glucose uptake, glycogen, glycolytic, and gluconeogenic flux rates were quantified using arteriovenous difference methods and [3-^3^H]-glucose tracer kinetics. Liver biopsies were collected before and after the afternoon clamp to assess gene transcription and protein regulators of hepatic glucose metabolism. During the afternoon clamp, despite matched insulin, glucagon, and glucose levels, net hepatic glucose uptake was 41% lower in AM INS+GCG (3.6±0.4 mg/kg/min) than in AM INS (6.1±0.6 mg/kg/min; p<0.003). This was accompanied by a trend toward incomplete suppression of hepatic glucose production in AM INS+GCG (1.4±0.4 mg/kg/min), whereas it was fully suppressed in the AM INS group (p=0.06). Direct glycogen synthesis was also 44% lower in AM INS+GCG (1.8±0.2 vs 3.2±0.7 mg/kg/min; p<0.015), along with reductions in net glycogen synthesis and glycolytic flux. Morning insulin with basal glucagon increased hepatic glucokinase mRNA and protein before the afternoon clamp, whereas concurrent glucagon prevented this induction. In summary, antecedent morning hyperglucagonemia attenuates insulin-mediated hepatic priming, reducing hepatic glucose flux during a later hyperinsulinemic-hyperglycemic challenge. These findings identify glucagon as a regulator of hepatic metabolic memory alongside insulin and demonstrate that early-day insulin-glucagon dynamics shape the liver’s response to subsequent challenges, providing mechanistic insight into postprandial glucose regulation and implications for metabolic health and diabetes risk.

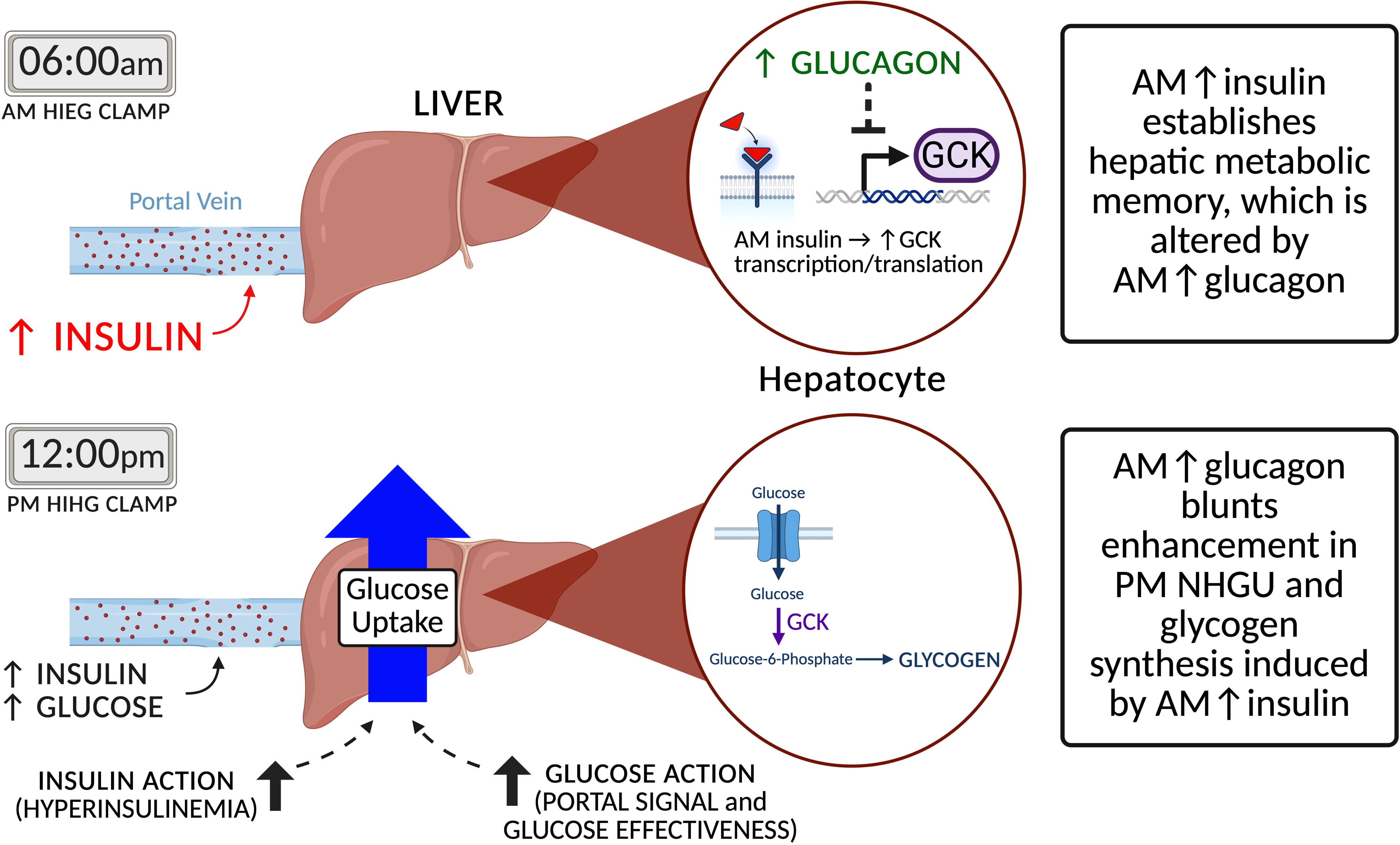

## 1 Introduction

The body’s metabolic response to the first meal of the day influences glucose regulation during subsequent meals, improving postprandial glucose handling during the second meal (1–4). We have shown that this response, known as the Staub-Traugott effect or second meal phenomenon, is a result of enhanced hepatic glucose uptake (HGU) and hepatic glycogen storage in the afternoon, processes that are altered by morning insulin exposure (5). While insulin is now recognized as a central driver of this response (5, 6), the roles of other hormonal and metabolic factors remain unclear.

Hepatic glucose metabolism operates within a broader systemic network, where signaling feedback loops with other tissues, mediated by hormones, nutrient flux, and neural pathways, help maintain whole-body energy balance (7–9). One key factor that may influence the second meal effect is glucagon, which is canonically known to acutely oppose insulin action in the liver by stimulating hepatic glucose production (HGP), primarily through glycogenolysis and, to a lesser extent, gluconeogenesis (10, 11). Since insulin and glucagon are typically co-secreted in response to mixed meals, their interaction is critical for shaping postprandial glucose metabolism throughout the day (12, 13). Previous studies have demonstrated that elevated glucagon levels can acutely reduce HGU and blunt glycogen synthesis during a single meal, even when insulin levels are high (14, 15). Notably, in individuals with prediabetes or type 2 diabetes, the postprandial glucagon response to mixed- or high-protein meals is often exaggerated compared to normoglycemic individuals, contributing to impaired glucose homeostasis and increased risk of disease progression (13, 16–18). Disruptions in systemic insulin or glucagon dynamics can therefore lead to widespread metabolic dysregulation. While glucagon’s acute effects are well characterized, its morning effects on glucose handling later in the day have never been investigated. These complex inter-meal regulatory networks require mechanistic studies that examine how the liver integrates both hormonal and metabolic signals across consecutive meals (19, 20).

Metabolic memory, which describes the ability of tissues to retain lasting effects from prior metabolic or nutritional exposures, is an emerging concept relevant to glucose regulation. Rather than fully resetting after each meal or physiological state, cells can carry forward molecular and functional imprints from past stimuli that shape subsequent responses. Most research to date has focused on long-term adaptations in tissues such as adipose and skeletal muscle, where chronic obesity, exercise, or metabolic stress produce durable molecular or functional changes (21, 22). For example, adipose tissue retains obesity-induced epigenetic marks even after weight loss, reflecting a memory of prior stress (23, 24). Similarly, skeletal muscle shows persistent adaptations following exercise or metabolic challenges, demonstrating functional memory beyond the initial exposure (25, 26). These findings indicate that metabolic responses are not merely acute but are conditioned by prior exposures, which alter tissue sensitivity and systemic nutrient handling over time. However, metabolic memory in the liver remains poorly understood under both acute and chronic conditions. It is well known that many hepatic processes, including those regulating glycogen storage and transcriptional changes, are transient and reversible, raising the question of how and for how long the liver can retain the effects of prior hormonal or nutritional signals (27).

We therefore aimed to determine whether a morning rise in glucagon alters the insulin-driven priming of afternoon net hepatic glucose uptake and glycogen storage. We hypothesized that morning hyperglucagonemia attenuates insulin’s effects on afternoon hepatic glucose metabolism, potentially reducing the benefits of insulin priming at breakfast. By directly assessing these effects, we sought to clarify the interaction between insulin and glucagon across meals and to determine how hormonal fluctuations shape hepatic glucose handling over time. This knowledge is important, not only for understanding the mechanisms underlying the second meal effect, but also for informing therapeutic strategies to be used in diabetes management, including dual- hormone insulin and glucagon pump systems or co-agonists designed to optimize glycemic control by leveraging the complementary actions of insulin and glucagon (28).

## 2 Research Design and Methods

### 2.1 Animal care and surgical procedures

A total of 29 adult mongrel dogs (16 males, 13 females) were included. Dogs were allocated to experimental groups to approximate a 1:1 male-to-female ratio. The sample size was based on preliminary and historical data. Power analysis using G*Power indicating that 8 animals per group would provide approximately ≥95% power to detect differences in afternoon net hepatic glucose uptake, the primary outcome, as well as molecular endpoints. An alpha level of 0.05 was used, and the effect size and variance estimates were derived from prior canine clamp studies. Power to detect potential sex differences was approximately ≥80%; however, sex comparisons were not a primary outcome in the design and were therefore considered exploratory. Consistent with this, exploratory analyses did not reveal significant sex-specific effects on hepatic glucose metabolism. The dog model was used because of its suitability for repeated catheterization of the portal and hepatic veins, enabling detailed *in vivo* hepatic flux analysis under controlled conditions. In addition, canine metabolism closely resembles human metabolism, and their genomes share a high degree of similarity and conserved function, further supporting translational relevance (29, 30). 16 dogs (*n*=8 per group; AM INS and AM INS+GCG) underwent the full protocol, including a 4h AM hyperinsulinemic-euglycemic clamp, 1.5h rest, and a 2.5h afternoon hyperinsulinemic-hyperglycemic clamp (mean weight 25.2±0.8 kg). Tissues from 13 additional dogs were used for hepatic molecular analyses only. Five dogs served as basal hepatic tissue controls, and four dogs per clamp group underwent only the AM clamp and rest period in order to assess molecular readouts prior to the afternoon clamp.

All procedures were approved by the Vanderbilt University Institutional Animal Care and Use Committee (IACUC) and Division of Animal Care (DAC). Dogs were sourced from a USDA-licensed vendor and housed in AAALAC-accredited facilities. Two weeks before each experiment, laparotomy under general anesthesia was performed to place hepatic artery and portal vein flow probes; catheters were inserted into the hepatic vein, portal vein, femoral artery, splenic and jejunal veins (for portal vein infusion), and inferior vena cava, with terminal ends secured subcutaneously until study day. Dogs were maintained on a once-daily controlled diet (46% carbohydrate, 34% protein, 14.5% fat, 5.5% fiber) and fasted for 18h before experiments. Only healthy animals were included, defined as consuming ≥75% of the prior meal, with leukocyte count <18,000/mm^3^ and hematocrit >34%. No animals were excluded. Blood sampling did not exceed 20% of total blood volume.

### 2.2 Experimental Design

The 8-hour protocol included two clamping periods designed to replicate postprandial pancreatic hormone profiles at times corresponding to breakfast (AM clamp, 0-240 min) and lunch (PM clamp, 330-480 min) **(Fig. 1).** Blood was collected every 15-30 min from the femoral artery, portal vein, and hepatic veins, while arterial glucose was monitored every 5 min, and peripheral glucose infusion was adjusted to maintain target glucose levels.

**Figure 1.**
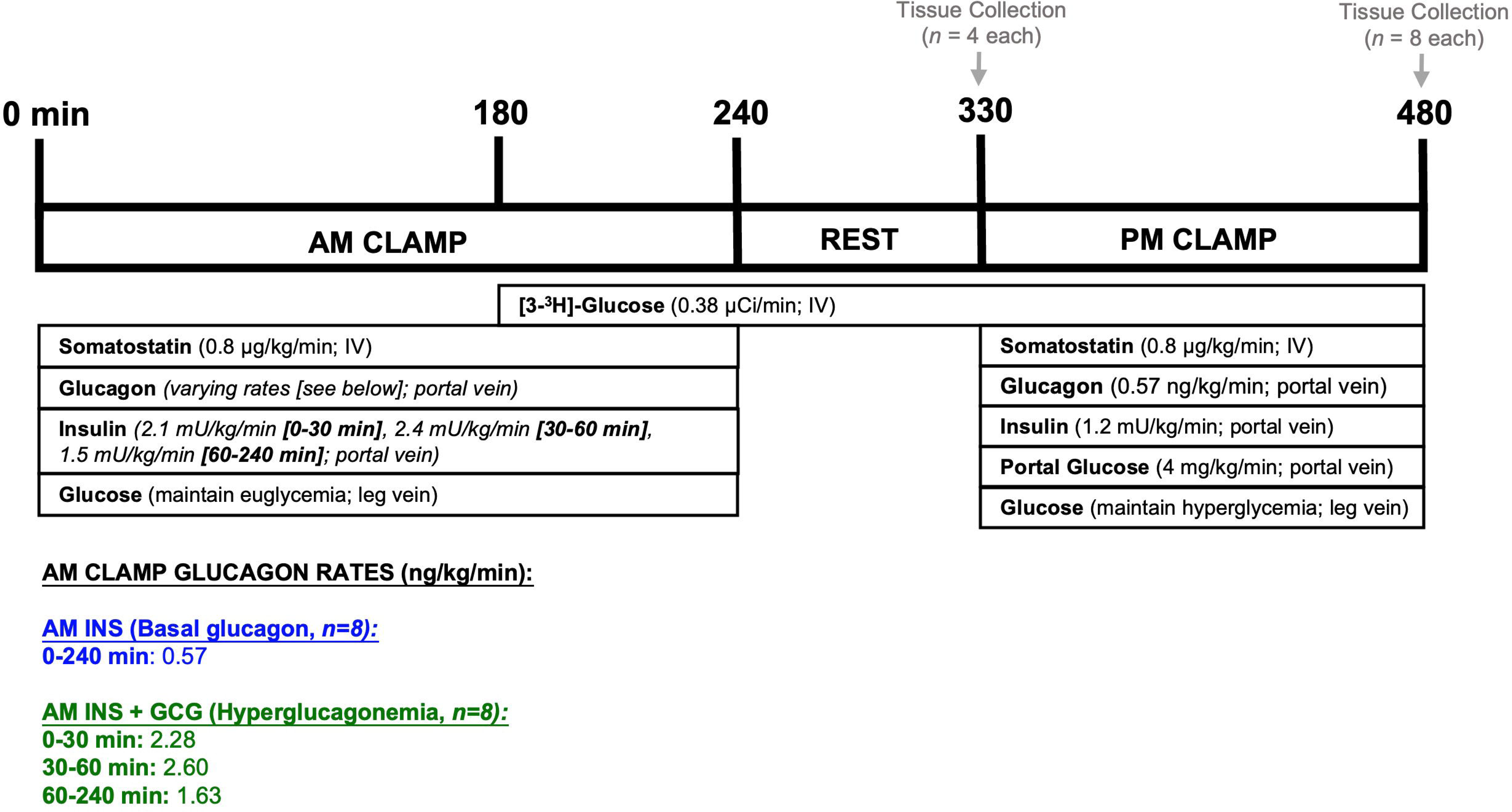
Experimental protocol. Canines underwent a 4h euglycemic clamp in the morning (AM; 0-240 min). Somatostatin was infused to suppress endogenous secretion of glucagon (GCG) and insulin (INS). Insulin was replaced in a pattern to mimic levels previously observed during a morning duodenal glucose infusion. One group received basal glucagon (AM INS), while the other received hyperglucagonemia (AM INS+GCG) in a pattern maintaining an equal molar ratio between insulin and glucagon throughout the AM clamp. A tracer infusion was initiated near the end of the AM clamp (180 min) to allow equilibration before the start of the PM clamp. Following the AM clamp, there was a 1.5h rest period (240-330 min) during which all infusions were halted. At the end of this period, a subset of dogs (n = 4/group) was euthanized for hepatic tissue collection and molecular analysis. The remaining dogs (n = 8/group) underwent a 2.5h PM HIHG clamp with portal glucose delivery (330-480 min), after which hepatic tissue was collected. Additional details are provided in the Methods section. AM INS, blue; AM INS+GCG, green.

#### 2.2.1 Morning (AM) Clamp Period (0-240 min)

We previously showed that a physiological morning insulin rise, rather than hyperglycemia, drives hepatic metabolic memory that enhances afternoon HGU (5). Accordingly, a hyperinsulinemic-euglycemic clamp was performed in the morning. At the start of the clamp, somatostatin (Bachem) was infused into the inferior vena cava (0.8 µg/kg/min) to suppress endogenous insulin and glucagon secretion. Intraportal insulin (Novolin R; Novo Nordisk) was infused at 2.1 mU/kg/min (0-30 min), 2.4 mU/kg/min (30-60 min), and 1.5 mU/kg/min (60-240 min) to replicate the physiologic insulin secretion pattern previously observed during a morning duodenal glucose infusion (5).

Two experimental groups were studied. One group (AM INS) received a basal intraportal glucagon infusion (GlucaGen; Novo Nordisk) at 0.57 ng/kg/min for 0-240 min, while the other group (AM INS+GCG) received an elevated, physiologic infusion of glucagon designed to maintain a consistent insulin-to-glucagon molar ratio over the AM clamp period (2.28 ng/kg/min, 0-30 min; 2.60 ng/kg/min, 30-60 min; 1.63 ng/kg/min, 60-240 min). These conditions were designed to mimic the morning hormonal profile following a carbohydrate- or mixed-meal, respectively. Glucose was clamped at euglycemia (∼100 mg/dL plasma glucose). To assess afternoon HGU and direct glycogen synthesis, a peripheral [3-^3^H]-glucose infusion (Revvitty) was initiated at 180 min (38 µCi prime, continuous 0.38 µCi/min infusion). At the end of the AM clamp (240 min), all infusions except the tracer were discontinued.

#### 2.2.2 Non-clamp Period (240 – 330 min)

To assess glucose kinetics before the PM clamp, blood was sampled from the femoral artery, portal vein, and hepatic veins at 300, 315, and 330 min **(Fig. 1).** A subset of dogs (*n*=4/group) were anesthetized at 330 min to examine the liver’s molecular profile at the start of the PM period. Hepatic tissue was rapidly harvested, flash-frozen in liquid nitrogen to preserve cellular conditions, and stored at −80°C, followed by euthanasia with pentobarbital.

#### 2.2.3 Afternoon (PM) Clamp Period (330 – 480 min)

Both AM INS and AM INS+GCG groups (*n*=8/group) underwent an identical 2.5h hyperinsulinemic-hyperglycemia (HIHG) clamp in the afternoon **(Fig. 1).** Somatostatin (0.8 µg/kg/min) was infused peripherally, while basal glucagon (0.57 ng/kg/min) and fourfold basal insulin (1.2 mU/kg/min) were administered intraportally. Portal glucose was infused at 4 mg/kg/min to activate the portal vein glucose signal (31), and a primed peripheral infusion raised plasma arterial glucose to hyperglycemic levels (∼200 mg/dL), maintained throughout the PM clamp. These conditions were designed to replicate a steady-state metabolic environment following a carbohydrate-rich afternoon “meal” (32). After the final sample (480 min), dogs were anesthetized, hepatic tissue was collected and preserved, and the dogs were euthanized as described above.

### 2.3 Analyses

#### 2.3.1 Biochemical and Molecular Methods

Whole blood samples were utilized to assess hormone and substrate flux across the liver using arteriovenous balance techniques (33). Plasma glucose levels were measured in quadruplicate (Analox GM9). Insulin (#PI-12K, MilliporeSigma), glucagon (#GL-32K, MilliporeSigma), and cortisol (VUMC Analytical Services in-house primary antibody with I^125^ cortisol from MP Biomedicals) were quantified via radioimmunoassay (33). Glucagon values were background corrected as described elsewhere (34) to account for nonspecific binding, ensuring that reported glucagon levels accurately reflect true circulating concentrations and strengthening the reliability of hormonal flux analyses. Key intermediates, including lactate, glycerol, alanine, and non-esterified fatty acids, were measured using enzymatic spectrophotometric assays (33). Plasma samples were deproteinized using the Somogyi method and quantified via liquid scintillation counting to determine the specific activity of [3-^3^H]-glucose in each sample (35).

Hepatic tissue collected at the start and end of the PM clamp was analyzed for glycogen content, gene expression, protein abundance, and enzyme activity using validated canine-specific methods (33, 36). Detailed protocols for reagents, primers, antibodies, and calculations are provided in the Supplementary Materials.

##### 2.3.1.1 RNA Extraction and Quantitative PCR

Total RNA was isolated from ∼50 mg of frozen liver using Tri-reagent (Sigma-Aldrich) and purified with the Direct-zol RNA Miniprep Kit (Zymo Research). RNA yield and purity were assessed spectrophotometrically (A260/A280>1.8). First-strand cDNA was synthesized from 1 µg RNA (High-Capacity cDNA Reverse Transcription Kit, Applied Biosystems). qPCR was performed on a CFX96 Real-Time PCR System (Bio-Rad) using SsoAdvanced Universal SYBR Green Supermix. Reactions contained 100 ng cDNA and 0.4 µM primers in 25 µL total volume. Cycling conditions: 95°C for 3 min, then 39 cycles of 95°C for 10 sec and 55°C for 30 sec. Melt curves confirmed specificity, and relative expression was calculated using the 2^–ΔΔCt method with GAPDH as reference. Primer details are listed in Supplemental Table 1.

##### 2.3.1.2 Western Blotting

Frozen liver (∼100 mg) was homogenized in 1 mL ice-cold buffer (20 mM Tris, 200 mM NaCl, 50 mM NaF, 1 mM EDTA, 1 mM EGTA, 10% glycerol, 1% SDS, pH 7.2) with protease and phosphatase inhibitors (Sigma-Aldrich). Homogenates were centrifuged at 3,000 × g for 10 min at 4°C. Protein concentration was determined by BCA assay (Bio-Rad) and adjusted to 4 µg/µL in Laemmli buffer. Samples (40 µg per lane) were separated on 4-12% Criterion TGX gels (Bio-Rad), transferred to nitrocellulose membranes, and stained with Ponceau S for total protein normalization. Membranes were blocked with 5% BSA in TBST and incubated overnight at 4°C with primary antibodies: pAkt (#9271, Cell Signaling), tAkt (#9272, Cell Signaling), pGS (#98348, Cell Signaling), tGS (#3893, Cell Signaling), pGP (#ab227043, Abcam), tGP (#ab198268, Abcam), pFOXO1 (#9461, Cell Signaling), tFOXO1 (#2880, Cell Signaling), and GK (#sc-17819, Santa Cruz). Primary antibody dilutions: 1:5,000 (GK 1:10,000; p/tFOXO1 1:1,000). After washing, membranes were incubated with 1:5,000 HRP-conjugated secondary antibodies for 1 h and developed with ECL reagents (GE Healthcare). Bands were quantified using ImageJ and normalized to basal controls.

##### 2.3.1.3 Hepatic Glycogen Content

Terminal liver glycogen content was measured using the amyloglucosidase method of Keppler and Decker (37). Briefly, frozen liver (∼180 mg) was homogenized in 0.6 N perchloric acid, neutralized with potassium bicarbonate, and incubated with amyloglucosidase (2 mg/mL, 40°C, 2h) to hydrolyze glycogen to glucose. Duplicate enzyme-free controls were processed in parallel. Glycogen content was calculated as glucose in enzyme-treated samples minus glucose in controls (oyster glycogen). Standards were processed alongside samples, and supernatants were analyzed for glucose concentration using a glucose analyzer (Analox GM9).

##### 2.3.1.4 Glucose-6-Phosphatase Activity Assay

Microsomal membranes were isolated from ∼1 g frozen liver (38). Tissue was homogenized in 50 mM HEPES, 25 mM KCl, 5 mM MgCl_2_, 0.25 M sucrose (pH 7.4), centrifuged at 7,800 × g for 6 min, and supernatant ultracentrifuged at 214,000 × g for 30 min. The microsomal pellet was resuspended in 29 mM MES, 21 mM Tris, 50 mM NaCl (pH 6.5). Microsomal protein (40 µg) was incubated with 10 mM G6P or M6P at 30°C for 15 min, reactions were quenched with 12% SDS, treated with Pi-chelating and developing solutions, and absorbance measured at 850 nm. Background Pi was subtracted, and activity expressed per mg microsomal protein per min.

##### 2.3.1.5 Glycogen Synthase Activity Assay

Glycogen synthase activity was measured by incorporation of UDP-[^14^C]-glucose into glycogen under low (active) and high (total) G6P conditions (2). Frozen liver (∼100 mg) was homogenized in potassium fluoride/EDTA/glycogen buffer (pH 7.0), centrifuged (500 × g, 10 min, 4°C), and supernatants diluted in pH 7.0 or 8.5 buffers. Aliquots were incubated with reaction buffer containing UDP-[^14^C]-glucose and low or high G6P. Samples were spotted on filter paper, precipitated in 70% ethanol, washed, air-dried, and counted by scintillation. Parallel blanks corrected for background incorporation, and activity was normalized to total protein.

#### 2.3.2 Calculations

Net hepatic glucose balance (NHGB) was determined using arterio-venous difference calculations, expressed as

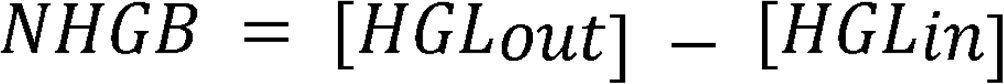

where HGLout represents the glucose output from the liver and was computed as

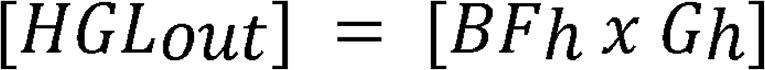

and HGLin denotes the glucose entering the liver, calculated as

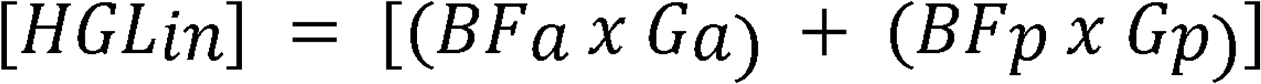

Here, G represents blood glucose concentration, BF indicates measured blood flow, and A, P, and H correspond to the hepatic artery, hepatic portal vein, and hepatic vein, respectively. Negative NHGB indicates net hepatic glucose uptake (NHGU), reflecting the liver’s contribution to whole-body glucose disposal (HGLin>HGLout). This methodology was extended to assess net hepatic balance for various substrates and hormones across the liver. Hepatic fractional glucose extraction (HFrEx) quantifies the proportion of glucose removed by the liver and was calculated as

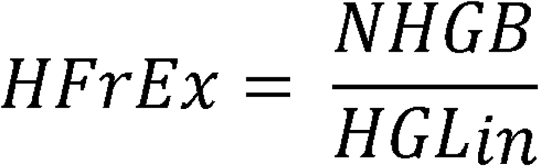

Tracer-determined unidirectional hepatic glucose uptake (HGU) was calculated as

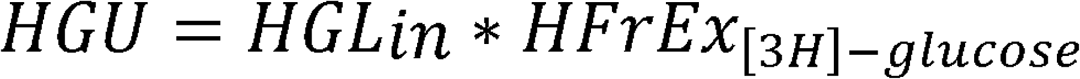

Hepatic glucose production (HGP) was calculated as the difference between NHGU and HGU. Hepatic sinusoidal substrate concentration (HSSC) was calculated as the flow-weighted average of arterial and portal vein concentrations

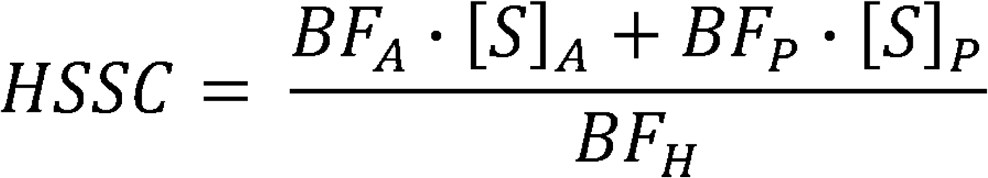

This represents the average substrate concentration in liver sinusoids, accounting for relative contributions from the hepatic artery and portal vein. Non-HGU, attributed primarily to skeletal muscle glucose uptake, was derived as

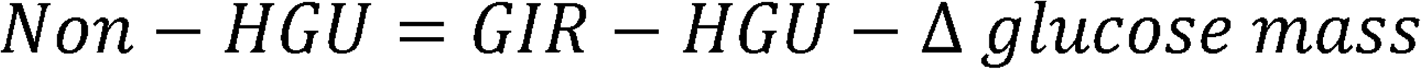

Where Δ glucose mass represents the change in glucose pool size between time points, accounting for extracellular distribution and pool fraction. The distribution volume was set to 22% of body weight with a pool fraction of 0.65 (39).

Direct glycogen synthesis, which quantifies glycogen synthesized from glucose taken up by the liver, was estimated as

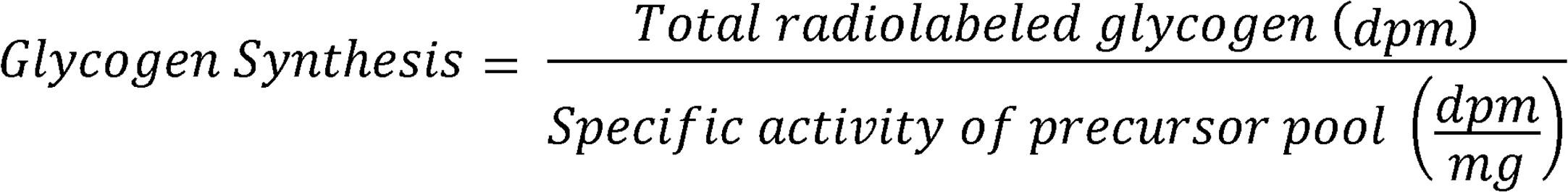

Where specific activity (SA) of the inflowing glucose precursor pool was calculated as

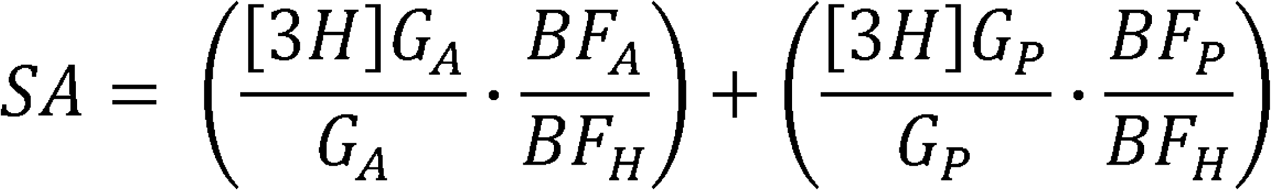

Where [3*H*]*G_A_* and [3*H*]*G_P_* represent the tracer concentrations in the arterial and portal blood supply, respectively. All gluconeogenic and glycolytic fluxes were expressed as glucose-equivalent units, reflecting conversion of three-carbon intermediates into six-carbon glucose equivalents. Total gluconeogenic flux was defined as conversion of gluconeogenic precursors (alanine and glycerol) to glucose-6-phosphate (G6P). Alanine uptake was doubled to account for total hepatic uptake of gluconeogenic amino acids, as alanine represents approximately 50% of this substrate pool (36, 40). Fluxes from alanine and glycerol were quantified as net hepatic alanine uptake (NHAU) and net hepatic glycerol uptake (NHGlyU), respectively. Therefore, gluconeogenic flux was calculated as

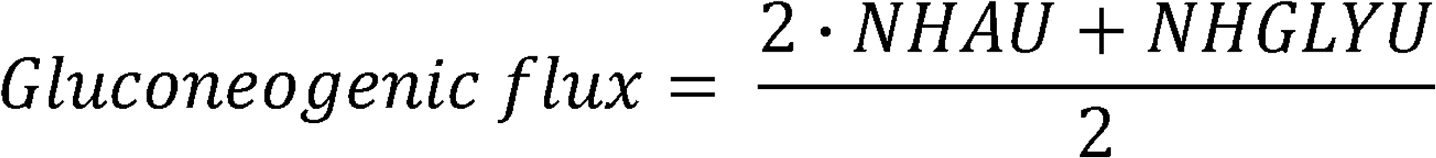

Glycolytic flux was estimated as

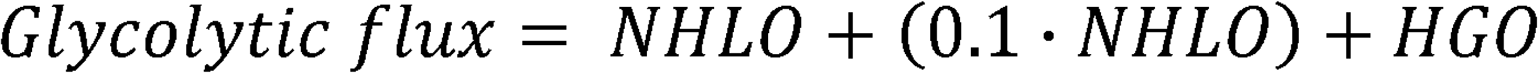

This represents the sum of net hepatic lactate output (NHLO), the estimated contribution of pyruvate, and hepatic glucose oxidation (HGO). NHLO was used as the primary measured lactate flux, with pyruvate production estimated as 10% of lactate flux (41), based on prior tracer studies demonstrating a stable lactate-pyruvate relationship across a wide range of clamp conditions. Hepatic glucose oxidation was assumed to be 0.2 mg/kg/min, consistent with prior canine clamp studies showing low and relatively invariant rates across insulin and glucose conditions (42, 43). Importantly, identical assumptions were applied across groups to ensure valid between-group comparisons.

Net hepatic gluconeogenic/glycolytic flux (NHGNG) was calculated as

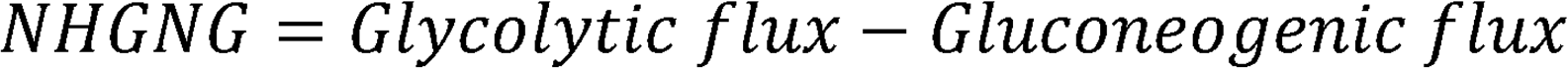

such that positive values indicate net glycolysis, and negative values indicate net gluconeogenesis. Net glycogen flux was then calculated to reflect net glycogen synthesis or breakdown as

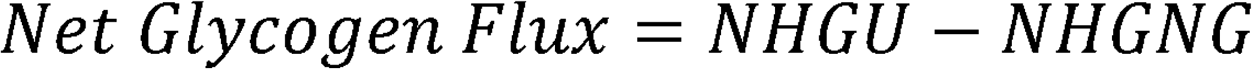

Where positive values indicate net glycogenesis and negative values indicate glycogenolysis. Together, these calculations provide integrated estimates of hepatic carbon partitioning between glucose production, glycolysis, and glycogen metabolism (30). All component fluxes were derived using previously validated assumptions, including estimates of pyruvate contribution, hepatic glucose oxidation, and the relative contribution of alanine to total gluconeogenic amino acid uptake, as described above.

#### 2.3.3 Statistics

Data are expressed as mean ± SEM. A two-way repeated-measures ANOVA was conducted to evaluate differences between groups and over time in flux analyses, followed by Tukey’s post hoc test for multiple comparisons. An unpaired two-tailed t-test was applied to analyze respective areas under the curve (AUC). ΔAUCs were calculated as the AUC relative to baseline. Molecular analyses were assessed using one-way ANOVA with Tukey’s post hoc test. Normality of all datasets were confirmed using the Shapiro-Wilk test, and all data were found to follow a normal distribution. Statistical analyses were performed using GraphPad Prism software. Exact p values are reported where available; otherwise, significance is indicated by threshold values (e.g., p <0.05, p <0.01, p <0.001, p <0.0001). A significance threshold of p <0.05 was applied.

### 2.4 Data availability

Values for all data points shown in figures are provided in the Supporting Data Values file. In addition, the dataset and supplementary materials are publicly available at doi.org/10.6084/m9.figshare.29988931. All other data generated and analyzed during the current study are available from the corresponding author upon reasonable request.

## 3 Results

### 3.1 AM Clamp Glucose and Hormone Data

During the AM clamp, euglycemia was successfully maintained in both groups **(Fig. 2A)**. Plasma insulin increased in a similar pattern across arterial, portal vein, and calculated hepatic sinusoidal sites in both the AM INS and the AM INS+GCG groups (**Fig. 2B-D Table 1).** Arterial plasma glucagon remained basal in the AM INS group (AM mean of 23 ± 1 pg/mL) but doubled in the AM INS+GCG group (AM mean of 49 ± 2 pg/mL; **Fig. 2E**, **Table 1).** Similar patterns were observed in portal vein and hepatic sinusoidal glucagon levels **(Fig. 2F, G**, **Table 1).** In the AM INS+GCG group, glucagon peaked at 60 min (arterial 71 ± 3 pg/mL; hepatic sinusoidal 116 ± 6 pg/mL) before gradually declining, yet it remained elevated relative to basal levels for the remainder of the clamp. Despite higher glucagon levels in the AM INS+GCG group, the glucose infusion rate required to maintain euglycemia was comparable between groups (mean of 9.5 ± 0.5 mg/kg/min in AM INS vs. 9.3 ± 1.3 mg/kg/min in AM INS + GCG, **Fig. 2H**). Net hepatic glucose balance switched from output to uptake in both groups, with no significant difference between the two groups **(Fig. 2I).**

**Figure 2.**
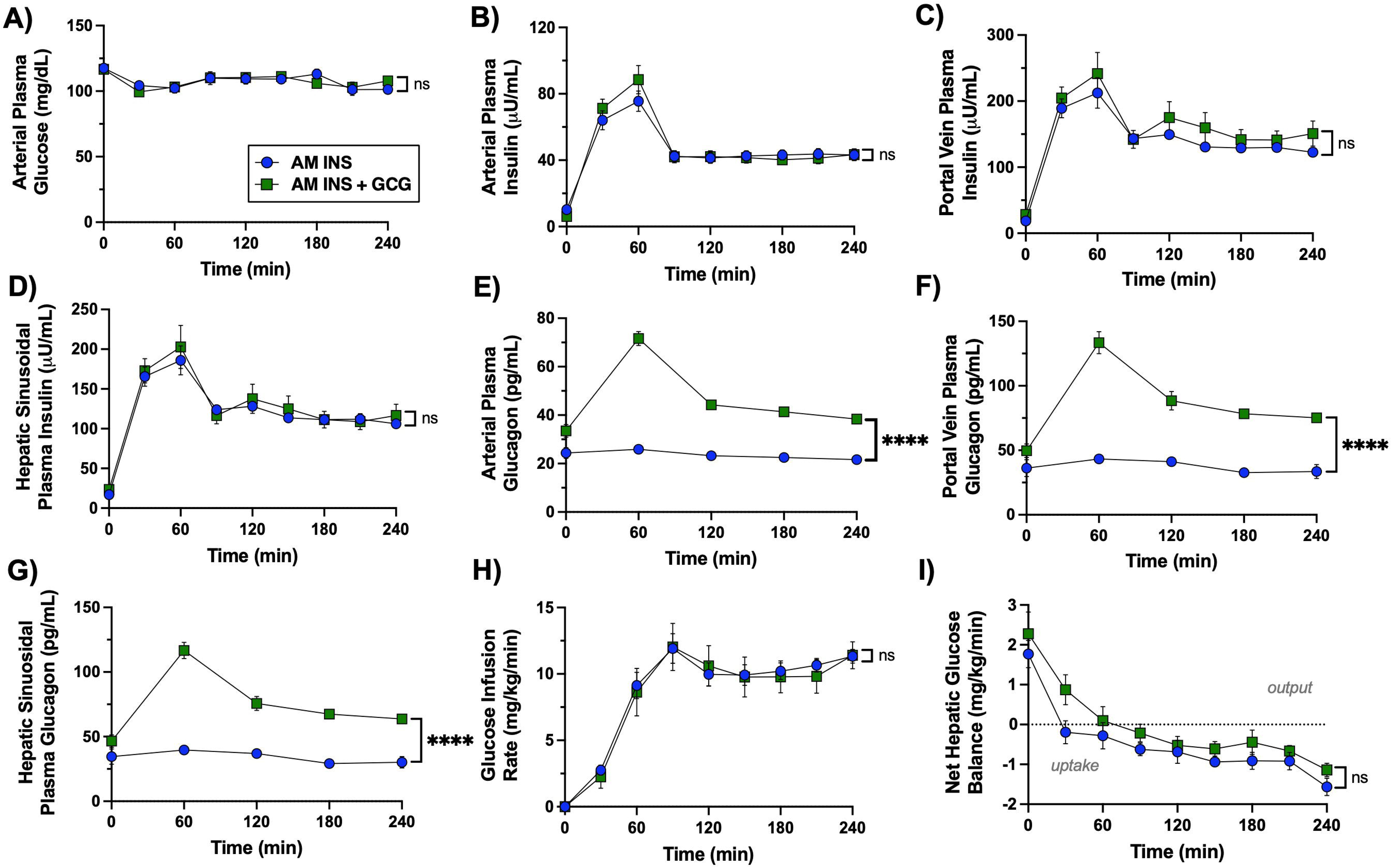
Morning (AM) clamp glucose and hormone flux data. (A) Arterial plasma glucose, (B) arterial plasma insulin, (C) portal vein plasma insulin, (D) hepatic sinusoidal plasma insulin, (E) arterial plasma glucagon, (F) portal vein plasma glucagon, (G) hepatic sinusoidal plasma glucagon, (H) glucose infusion rate, and (I) net hepatic glucose balance (negative values indicate uptake, positive values indicate output) are shown for the AM INS and AM INS+GCG groups (n = 8/group, except for F and G which have n = 4 for AM INS, and I which has n = 6 for AM INS) throughout the AM clamp. Data are expressed as mean ± SEM. ***P < 0.001, ****P < 0.0001 between groups; ns = non-significant. AM INS, blue; AM INS+GCG, green.

**Table 1.** Plasma hormone concentrations measured during the AM clamp.

| AM Clamp Parameter | Group |  |
| --- | --- | --- |
|  | AM INS (*n*=8) | AM INS+GCG (*n*=8) |
| Arterial Plasma Insulin<br>(μU/mL) | 50 ± 4 | 51 ± 4 |
| Hepatic Sinusoidal Plasma<br>Insulin (μU/mL) | 151 ± 6 | 170 ± 15 |
| Arterial Plasma Glucagon<br>(pg/mL) | 23 ± 1 | 49 ± 2**** |
| Hepatic Sinusoidal Plasma<br>Glucagon (pg/mL) | 34 ± 3 | 81 ± 3**** |

Data are presented as mean ± SEM. Insulin samples were analyzed every 30 min, and glucagon samples were analyzed every hour during the experimental protocol. Each value represents the mean of all samples collected within each clamping period (*n* = 8/group). \*\*\*\**P* < 0.0001.

After the 1.5h rest period, hormone and substrate levels had returned toward baseline, and all dogs were in a glucose-producing state **(Figs. 3-5)**. This reflects normalization of glucose homeostasis following the AM intervention and is evident across the 300-330 min period preceding the onset of the PM clamp.

**Figure 3.**
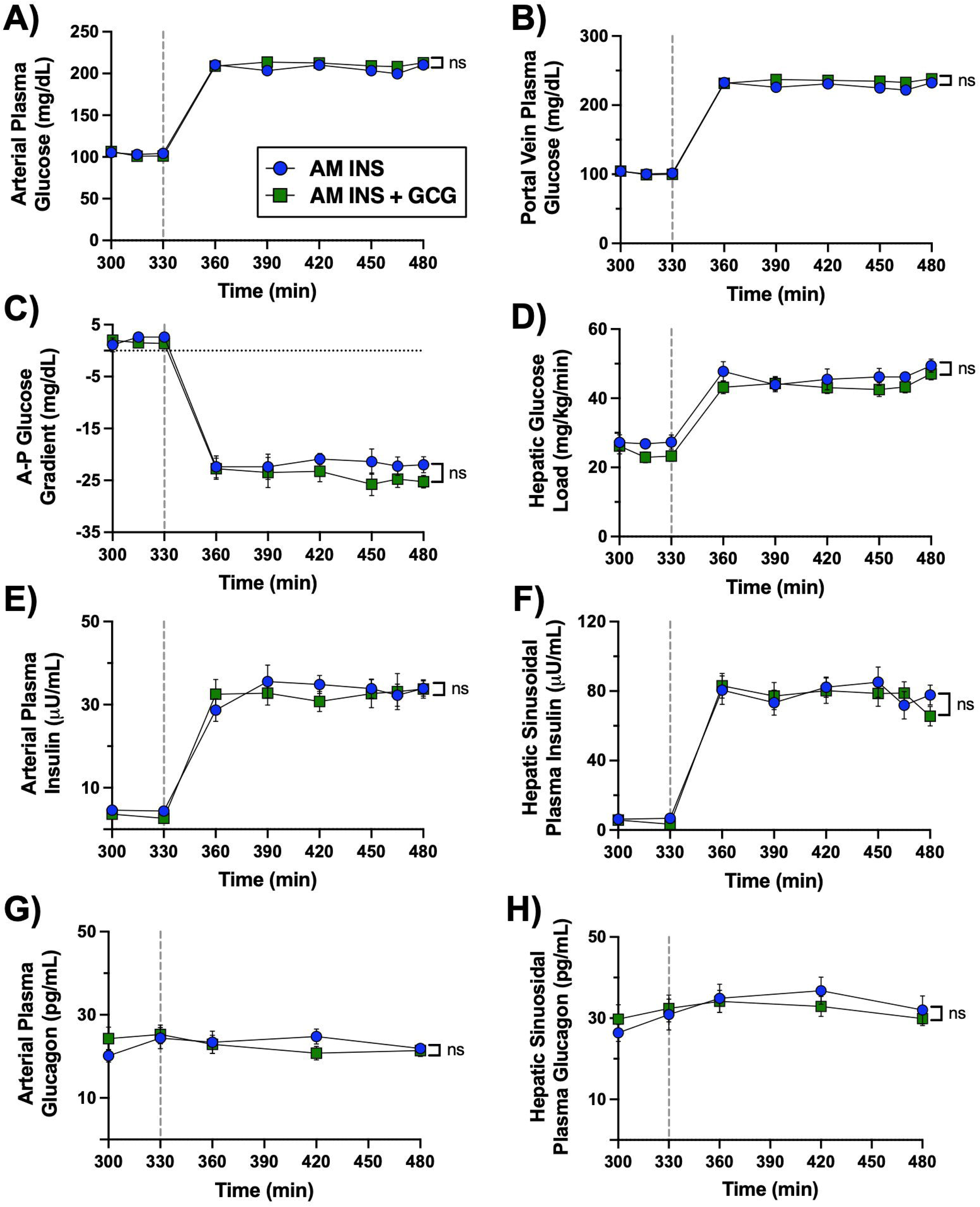
Afternoon (PM) clamp glucose and hormone flux data. A vertical line at 330 min indicates the end of the rest period and the onset of the PM clamp. (A) Arterial plasma glucose, (B) portal vein plasma glucose, (C) arterial–portal glucose difference, (D) hepatic glucose load, (E) arterial plasma insulin, (F) hepatic sinusoidal plasma insulin, (G) arterial plasma glucagon, and (H) hepatic sinusoidal plasma glucagon are shown throughout the 2.5h hyperinsulinemic-hyperglycemic PM clamp in both the AM INS and AM INS+GCG groups (n = 8/group). No significant differences were observed between groups for any measured parameter. Data are expressed as mean ± SEM; ns = non-significant. AM INS, blue; AM INS+GCG, green.

### 3.2 PM Clamp Glucose and Hormone Data

The PM clamp was designed to mimic a postprandial hormonal and substrate milieu by maintaining hyperglycemia through peripheral (leg vein) glucose infusion, while concurrent portal vein glucose infusion established a physiologic meal-associated arterial-to-portal vein glucose gradient of ∼20-25 mg/dL **(Fig. 3A-D).** Insulin levels were markedly elevated in both the arterial circulation and hepatic sinusoids (**Fig. 3E, F)**, whereas glucagon levels remained basal throughout the PM clamp **(Fig. 3G, H).** Thus, during the PM clamp matching hormonal and glycemic conditions existed in both the AM INS and AM INS+GCG groups.

The glucose infusion rate required to maintain hyperglycemia was similar in both groups during the PM clamp **(Fig. 4A, B).** Mean non-HGU was 7.6 ± 0.9 mg/kg/min in the AM INS group and 9.4 ± 1.3 mg/kg/min in the AM INS+GCG group, with no significant difference between the two groups **(Fig. 4C, D).** NHGU in the PM was 41% lower in the AM INS+GCG group (PM mean of 6.1 ± 0.6 vs. 3.6 ± 0.4 mg/kg/min in the AM INS vs. AM INS+GCG groups, respectively, P < 0.0027 **Fig. 4E, F).** This reduction occurred in conjunction with a non-significant difference in unidirectional HGU in the PM (mean of 5.7 ± 0.5 mg/kg/min vs. 5.0 ± 0.3 mg/kg/min in the AM INS vs. AM INS+GCG group, respectively; **Fig. 4G, H).** In contrast, HGP was fully inhibited in the AM INS group but averaged 1.4 ± 0.4 mg/kg/min in the AM INS+GCG group during the PM clamp. Although the HGP flux showed only a trend toward significance (P = 0.06, **Fig. 4I**), the ΔAUC was significantly higher in the AM INS+GCG group compared to AM INS **(Fig. 4J).** Consistent with this, net glycolytic flux was significantly lower in the AM INS+GCG group (P < 0.024; **Fig. 4K)**, and net glycogenic flux was also reduced by 41% (P < 0.004; **Fig. 4L)**. Together, these data demonstrate that prior morning hepatic glucagon exposure reduced the ability of morning insulin to enhance hepatic glucose uptake and storage in the PM.

**Figure 4.**
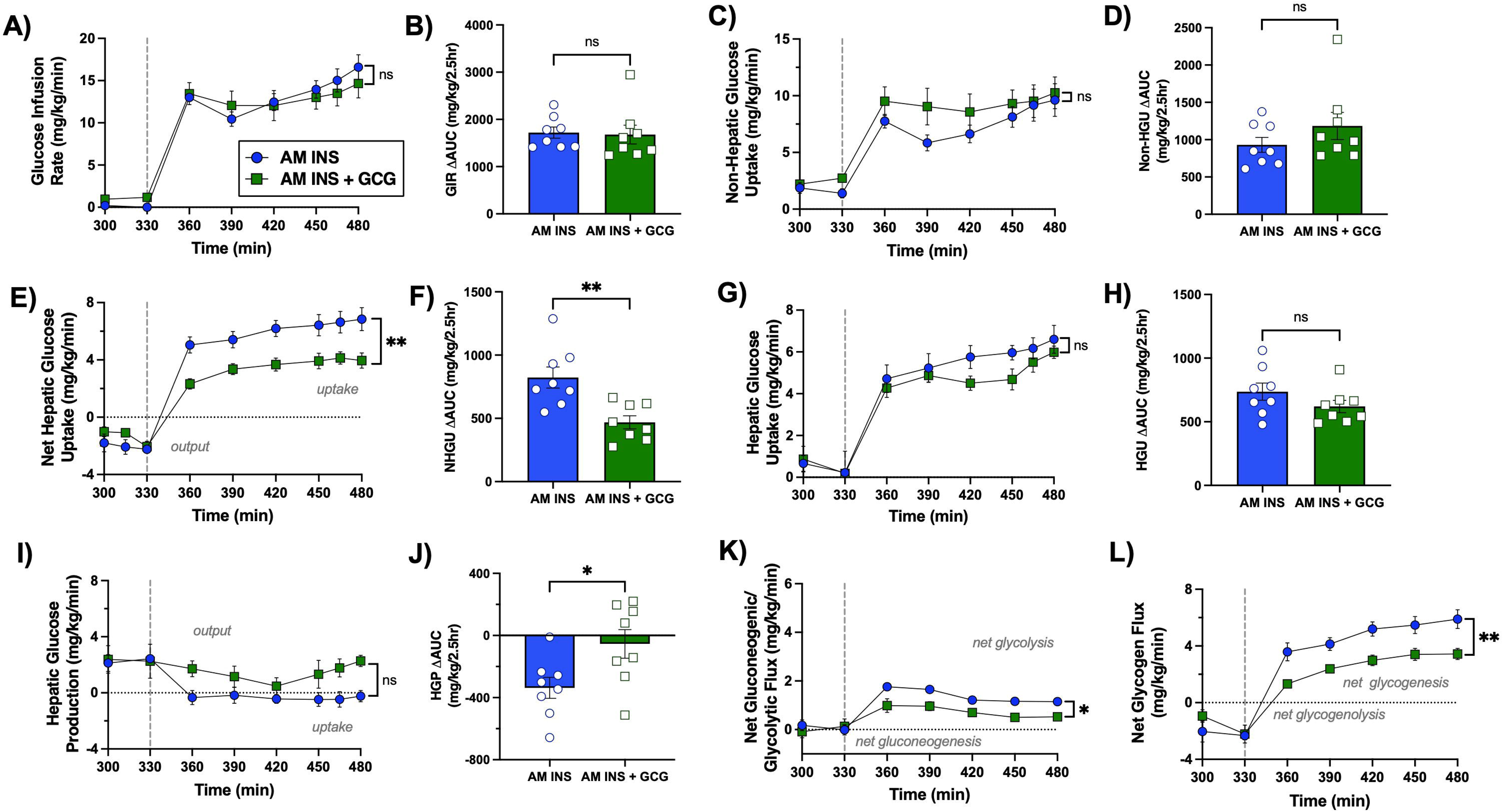
Glucose uptake and storage during the PM hyperinsulinemic-hyperglycemic clamp. A vertical line at 330 min indicates the end of the rest period and the onset of the PM clamp. (A) Glucose infusion rate (GIR), (B) GIR ΔAUC, (C) non-hepatic glucose uptake, (D) non-hepatic glucose uptake ΔAUC, (E) net hepatic glucose uptake (NHGU; values >0 indicate uptake, <0 indicate output), (F) NHGU ΔAUC, (G) unidirectional hepatic glucose uptake determined using tritiated glucose, (H) HGU AUC, (I) hepatic glucose production (HGP; determined from the difference between NHGU and HGU), (J) HGP AUC, (K) net gluconeogenic/glycolytic flux (values >0 indicate net glycolysis, <0 indicate net gluconeogenesis), and (L) net glycogen flux (values >0 indicate net glycogenesis, <0 indicate net glycogenolysis) are shown over time for the AM INS and AM INS+GCG groups (n = 8/group). Data are expressed as mean ± SEM. *P < 0.05, **P < 0.01 between groups; ns = non-significant. AM INS, blue; AM INS+GCG, green.

During the PM clamp, arterial plasma non-esterified fatty acid (NEFA) and glycerol levels, as well as net hepatic uptake of both substrates, were similarly suppressed in the AM INS and AM INS+GCG groups reflecting effective insulin-mediated inhibition of hepatic fatty acid handling under PM HIHG conditions **(Fig. 5A-D).** Circulating alanine levels and hepatic alanine balance remained stable throughout the PM clamp, with no differences between groups **(Fig. 5E, F)**. In contrast, arterial lactate concentrations and net hepatic lactate output were significantly lower in the AM INS+GCG group compared with the AM INS group (**Fig. 5G, H)**. These results indicate that antecedent morning glucagon exposure selectively reduced net hepatic glucose uptake and its downstream utilization in glycogen synthesis and glycolytic pathways during the PM clamp, while lipid and amino acid metabolism remained unaffected.

**Figure 5:**
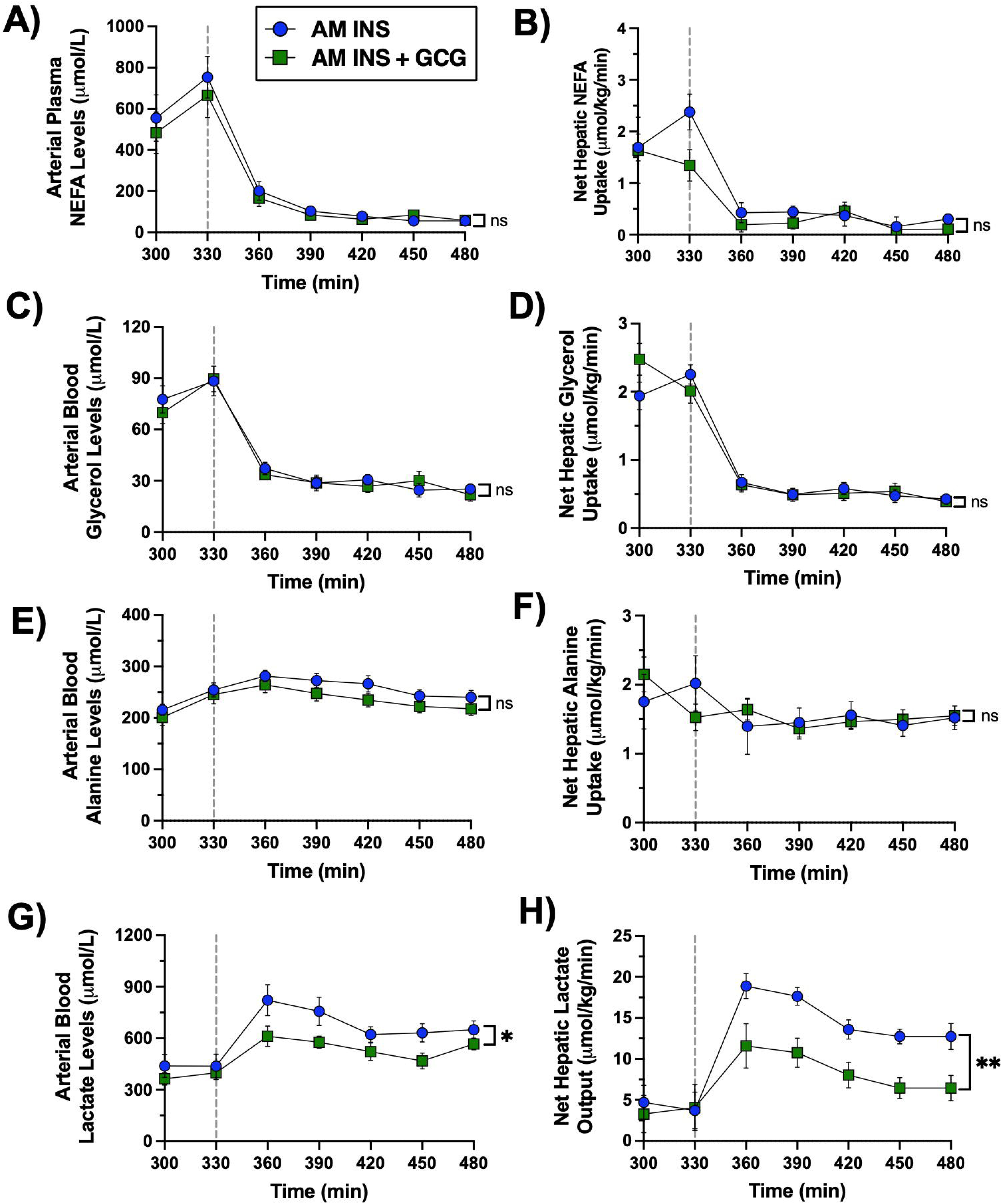
PM clamp fatty acid and metabolite flux data. A vertical line at 330 min separates the resting period from the onset of the PM clamp. (A) Arterial plasma non-esterified fatty acids (NEFA), (B) net hepatic NEFA uptake, (C) arterial blood glycerol, (D) net hepatic glycerol uptake, (E) arterial blood alanine, (F) net hepatic alanine uptake, (G) arterial blood lactate, and (H) net hepatic lactate output are shown throughout the PM HIHG clamp for the AM INS and AM INS+GCG groups (n = 8/group). Data are expressed as mean ± SEM. *P < 0.05, **P < 0.01 between groups; ns = non-significant. AM INS, blue; AM INS+GCG, green.

### 3.3 Terminal Liver Biopsies

Terminal liver biopsies were first obtained at 330 min, following completion of the AM clamp and rest period, but prior to initiation of the PM clamp **(Fig. 6A).** The 330 min time point was chosen to provide a baseline assessment of hepatic mRNA, protein, and metabolite levels prior to the PM clamp. Glucose flux measurements diverge between groups immediately upon initiation of the PM clamp, indicating that hepatic molecular changes had already occurred by that time. At 330 min, no differences in Akt phosphorylation were observed among groups **(Fig. 6B).** Glucokinase (*GCK*) mRNA was significantly increased in the AM INS group compared with basal, whereas this induction was absent in AM INS+GCG **(Fig. 6C).** Consistent with transcript levels, GCK protein expression was elevated only in AM INS **(Fig. 6D).** Total glucose-6-phosphatase (G6Pase) activity in intact and disrupted microsomes, measured using G6P as a substrate, was not different between the AM INS and AM INS+GCG groups at this time point **(Fig. 6E).** G6Pase activity in disrupted microsomes, measured using M6P as a substrate, also showed no difference (data not shown). Phosphorylation of glycogen synthase (GS) did not differ between groups, either **(Fig. 6F).** Furthermore, phosphorylation and activity of glycogen phosphorylase (GP) were not significantly different between groups **(Fig. 6G, H).**

**Figure 6.**
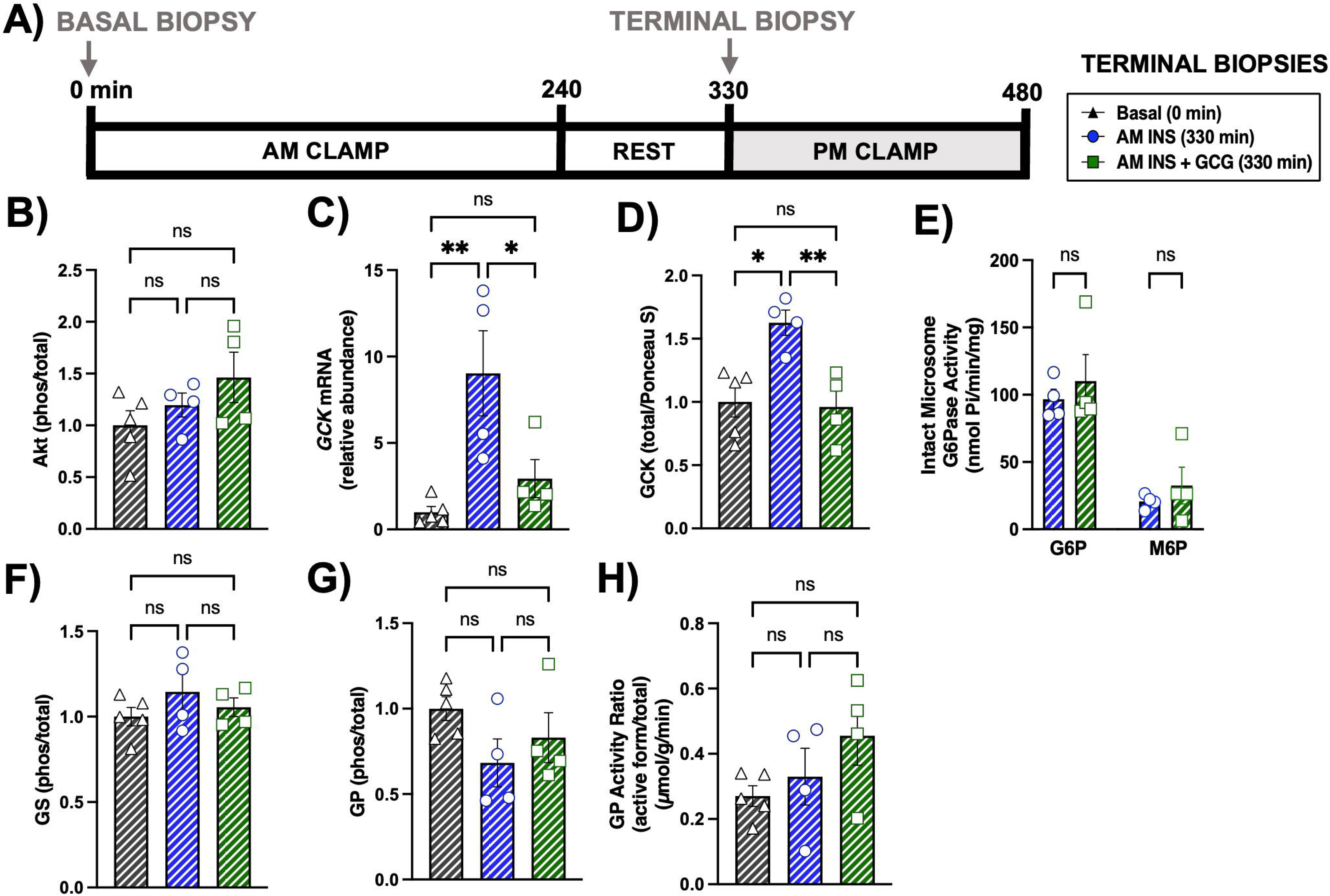
Terminal hepatic tissue transcript and protein analysis prior to the PM clamp. (A) Experimental timeline for hepatic biopsies collected at 0 minutes (Basal) and 330 minutes (terminal, immediately before the PM clamp for each experimental group). Levels of Akt protein (B), glucokinase mRNA (C), glucokinase protein (D), intact microsome glucose-6-phosphatase activity using G6P and M6P substrates (E), glycogen synthase protein (F), glycogen phosphorylase protein (G), and glycogen phosphorylase activity (H) are shown. Data are presented for the Basal (n = 5), AM INS (n = 4), and AM INS+GCG (n = 4) groups as mean ± SEM. *P < 0.05, **P < 0.01 between groups; ns = non-significant. Protein levels were measured by western blot and mRNA levels by real-time qPCR. Protein levels were normalized to Ponceau S total protein; mRNA levels were normalized to GAPDH. AM INS, blue; AM INS+GCG, green.

A separate set of biopsies was collected at 480 min, at the conclusion of the PM clamp **(Fig. 7A).** Akt phosphorylation was significantly increased in AM INS and AM INS+GCG compared with basal controls **(Fig. 7B).** *GCK* mRNA expression was induced in both groups compared to baseline **(Fig. 7C).** GCK protein remained significantly elevated only in AM INS, while levels in AM INS+GCG were not different from basal controls **(Fig. 7D).** Total glucose-6-phosphatase activity and activity in intact microsomes remained similar between the AM INS and AM INS+GCG groups **(Fig. 7E).** These measurements were performed in a subset of 4 animals per group. GS phosphorylation was comparably reduced in both experimental groups **(Fig. 7F),** accompanied by increased GS activity **(Fig. 7G).** GP phosphorylation was reduced in both groups compared with basal controls **(Fig. 7H),** while GP activity was not significantly different between any of the groups **(Fig. 7I).** Additionally, mean direct glycogen synthesis during the PM clamp (360-480 min) was significantly reduced in the AM INS+GCG group compared to the AM INS group (1.8 ± 0.2 vs. 3.2 ± 0.7 mg/kg/min, respectively, P < 0.015; **Fig. 7J).**

**Figure 7.**
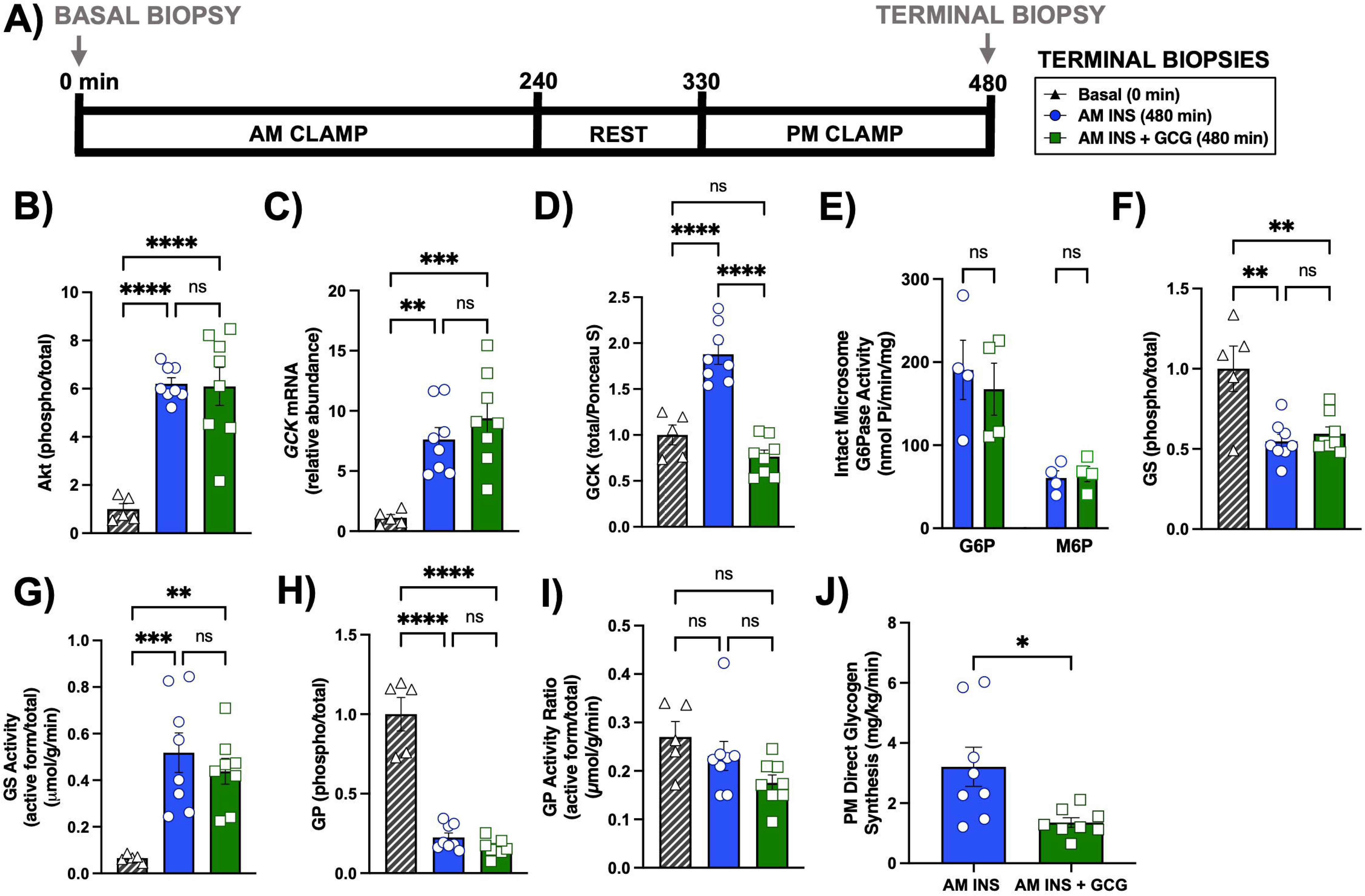
Terminal hepatic tissue transcript and protein analysis after the PM clamp. (A) Experimental timeline for hepatic biopsies collected at 0 minutes (Basal) and 480 minutes (terminal, end of the PM clamp for each experimental group). Levels of Akt protein (B), glucokinase mRNA (C), glucokinase protein (D), intact microsome glucose-6-phosphatase activity using G6P and M6P substrates for a subset of animals (n = 4/group; Basal not included; E), glycogen synthase protein (F), glycogen synthase activity (active/total; G), glycogen phosphorylase protein (H), glycogen phosphorylase activity (I), and mean direct glycogen synthesis during the PM clamp (J) are shown. Data are presented for the Basal (n = 5), AM INS (n = 8), and AM INS+GCG (n = 8) groups as mean ± SEM. *P < 0.05, **P < 0.01, ***P < 0.001, ****P < 0.0001 between groups; ns = non-significant. Protein levels were measured by western blot and mRNA levels by real-time qPCR. Protein levels were normalized to Ponceau S total protein; mRNA levels were normalized to GAPDH. AM INS, blue; AM INS+GCG, green.

In summary, AM insulin primed the liver by increasing *GCK* mRNA and protein prior to the PM clamp, whereas concurrent AM glucagon blocked this insulin-induced effect on *GCK* transcription and translation. At the end of the PM clamp, downstream regulation of glycogen synthase and phosphorylase were similar between experimental groups. While *GCK* mRNA levels were comparable at the end of the PM clamp, only the AM INS group maintained elevated GCK protein, which could partially explain the reduction observed in NHGU, glycolysis, and lactate release in the AM INS+GCG group. Supporting molecular data are shown in Supplementary Figures 1 and 2.

#### 3.4 Summary of Hepatic Fluxes

Figure 8 summarizes an integrated model of mean hepatic glucose fluxes during the PM clamp. In the AM INS group, NHGU was relatively large, reflecting sustained hepatic glucose retention following morning insulin exposure. This was entirely due to increased HGU, as HGP remained fully suppressed. Incoming glucose was phosphorylated to G6P, which was either stored as glycogen or metabolized through glycolysis and released as lactate or oxidized. In contrast, when elevated glucagon accompanied insulin in the morning, PM NHGU was lower, driven by a combination of a non-significant decrease in HGU and sustained HGP relative to the AM INS group. Additionally, less G6P was partitioned into glycolytic and glycogenic pathways. Together, these combined effects produced substantially lower hepatic glucose retention. These findings demonstrate that antecedent glucagon exposure induces metabolic memory, blunting insulin-mediated stimulation of NHGU and glycogen synthesis.

**Figure 8.**
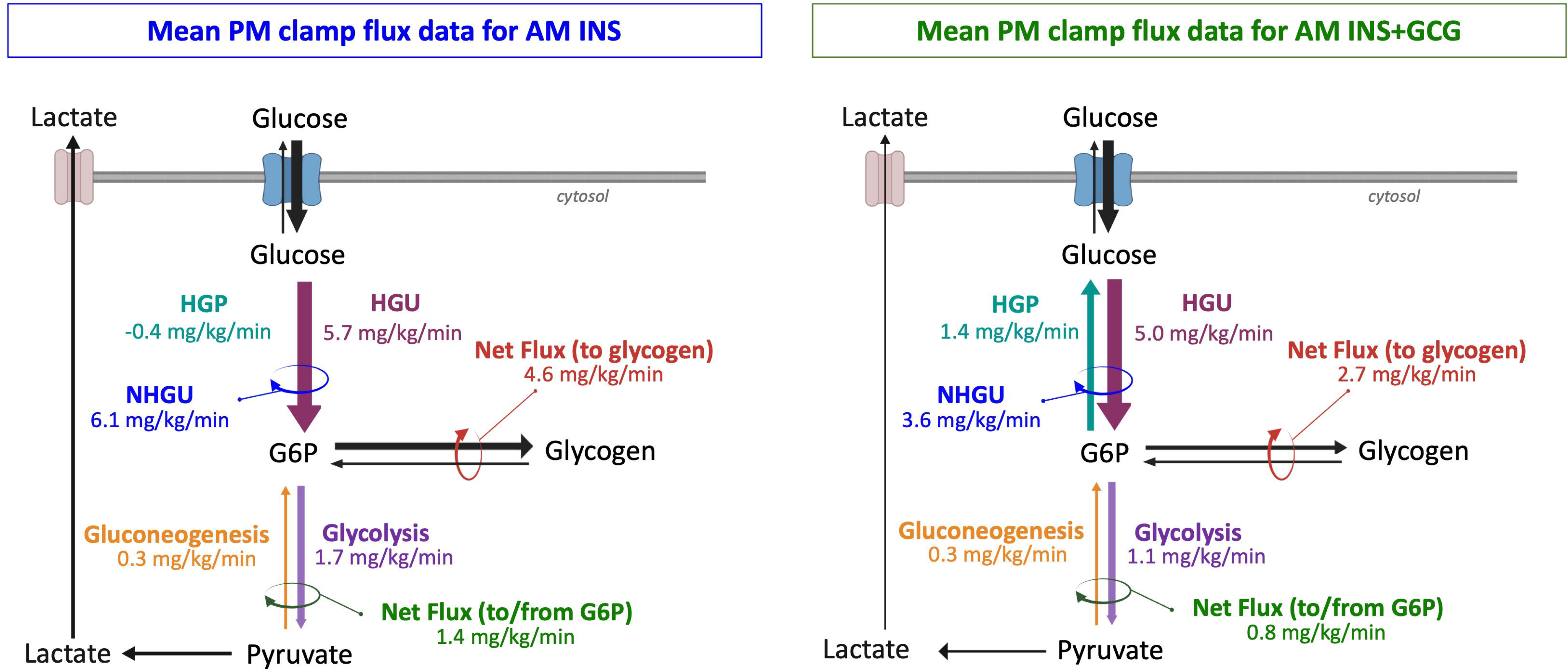
Summary of mean hepatic fluxes during the PM HIHG clamp. The left panel shows the AM INS group (n = 8), and the right panel shows the AM INS+GCG group (n = 8). Values represent mean fluxes ± SEM from 360-480 min of the PM clamp, determined by carbon balance analysis. Depicted fluxes include glucose to/from glucose-6-phosphate (G6P), G6P to/from glycogen, and G6P to/from pyruvate through gluconeogenesis and glycolysis (estimated using alanine, glycerol, and lactate as 3-carbon precursors). The magnitude of the arrows reflects the relative flow in each pathway. This schematic is integrative, based on calculated mean fluxes, and does not represent direct measurement of intracellular compartmental fluxes. Abbreviations: NHGU, net hepatic glucose uptake; HGU, hepatic glucose uptake; HGP, hepatic glucose production; Net HGU, net hepatic glucose uptake (HGU and HGP).

## 4 Discussion

Effective regulation of hepatic glucose metabolism is central to maintaining systemic glucose homeostasis. Breakfast plays a pivotal role in postprandial glycemic control and glycogen storage, influencing the metabolic response to subsequent meals in both healthy individuals and those with diabetes (1, 2, 44, 45). Skipping breakfast has been associated with an increased risk of developing type 2 diabetes, suggesting that disruptions in morning nutrient and hormone signaling can have persistent effects on glucose metabolism and overall metabolic health (46). Although the precise contribution of glucagon to metabolic dysregulation remains incompletely understood, postprandial hyperglucagonemia is consistently observed in obesity and type 2 diabetes, potentially impairing glucose homeostasis by limiting insulin-mediated glucose uptake and sustaining hepatic glucose production (13, 47). Since glucagon opposes insulin’s anabolic actions on the liver by promoting glycogenolysis and glucose output, understanding its impact on postprandial metabolism is particularly important given that the insulin-to-glucagon ratio governs hepatic metabolic balance (48). These considerations prompted us to investigate whether elevated morning glucagon impairs insulin-mediated priming of the liver for enhanced glucose uptake in the afternoon.

Our findings are among the first to demonstrate that antecedent morning glucagon exposure reprograms hepatic glucose metabolism thereby impacting NHGU and glycogen deposition later in the day. This highlights the sensitivity of the liver to changes in insulin and glucagon occurring hours before a second meal and underscores their importance to subsequent intracellular substrate partitioning and enzymatic regulation. These findings indicate that, like insulin, prior glucagon exposure establishes hepatic metabolic memory. Following morning glucagon exposure, afternoon carbon flux shifts away from storage and toward re-release, thereby attenuating insulin’s contribution to the second-meal effect. Notably, non-hepatic glucose uptake was unchanged across groups, indicating that these flux changes were specific to hepatic glucose metabolism and did not alter peripheral glucose disposal. Although glucose infusion rates were similar between groups despite differences in NHGU, the corresponding difference in non-hepatic glucose uptake did not reach statistical significance. As such, we cannot exclude the possibility of a modest effect of antecedent glucagon on peripheral glucose disposal; however, the lack of significance, together with the inherent variability in whole-body glucose utilization under these conditions, suggests that the primary effect was at the liver. Future studies will be necessary to specifically evaluate potential subtle effects in non-hepatic tissues that were not resolved in the present study design.

To understand the molecular mechanisms underlying our findings, hepatic *GCK* mRNA and protein levels were measured over the course of the study (basal, prior to, and at the end of the PM period). As expected from the known effects of insulin on *GCK* transcription, 5.5 hours after the start of the insulin prime, hyperinsulinemia increased *GCK* mRNA and protein in the AM INS group. Glucagon, however, blocked the transcriptional and translational effects of insulin infusing during the morning period. This is consistent with prior *in vitro* work done in cultured rat hepatocytes, in which glucagon rapidly repressed insulin-stimulated *GCK* transcription via a cAMP-dependent pathway under supramaximal insulin conditions, reducing mRNA accumulation within 30-45 minutes (49). Furthermore, in hepatocytes in which *GCK* transcription had already been induced by prolonged insulin exposure, glucagon (or a cAMP analog) also accelerated degradation of existing *GCK* mRNA, with an apparent half-life of ∼45 minutes (49).

During the PM clamp, hyperinsulinemia in the AM INS group maintained the equilibrium of *GCK* mRNA transcription and degradation that was established during the morning period, such that *GCK* mRNA remained similar at the 330 min and 480 min time points. As a result, GCK protein levels also remained elevated in the AM INS group. This suggests that the insulin-dependent homeostatic setpoint for *GCK* mRNA and protein levels was similar during the two periods. In contrast, at the end of the PM clamp, *GCK* mRNA levels, but not protein, was elevated in the AM INS + GCG group. This was to be expected, however, since we have previously shown that it typically requires two to four hours to show a measurable increase in GCK protein following mRNA induction by insulin (50). Thus, the 2.5h hyperinsulinemic PM period was long enough to increase *GCK* mRNA but not protein levels. Notably, although total hepatic GCK protein levels at the end of the PM clamp were similar between the basal and AM INS+GCG groups, it is important to consider subcellular localization when interpreting these findings. Glucokinase regulatory protein dynamically modulates GCK by sequestering it in the nucleus during low-glucose conditions and releasing it into the cytoplasm when glucose is abundant (51–53). Therefore, it is plausible that a larger fraction of GCK in the AM INS+GCG group resides in the cytosolic, active compartment, which could help explain why HGU remains elevated in the AM INS+GCG group despite similar total GCK protein levels to basal tissue. This highlights that total protein measurements alone may not fully capture the functional state of GCK *in vivo*. This highlights that the molecular processes by which morning glucagon disrupts insulin-driven GCK accumulation and hepatic glucose flux are not fully understood and warrant further investigation. The use of complementary models that allow more detailed mechanistic exploration beyond what is feasible in the canine model will be important to clarify these processes (52).

Several molecular mechanisms likely contributed to the reduced PM NHGU observed in the AM INS+GCG group. Central among these is glucokinase, whose activity directly correlates with hepatic glucose uptake (45). As noted above, the morning rise in glucagon in the AM INS+GCG group attenuated insulin’s effect on hepatic *GCK* mRNA and protein later in the day. This likely reduced glucose phosphorylation and, in turn, potentially diminished hepatic G6P availability in the AM INS+GCG group thereby limiting substrate availability for glycogen synthesis and glycolysis (51). Downstream enzymes, including GS and GP, showed no differences in dephosphorylation or intrinsic activity; however, a reduced G6P pool in the AM INS+GCG group would be expected to limit allosteric activation of GS and relieve suppression of GP activity, effects not fully captured by protein abundance or activity assays (54). Additionally, HGP remained incompletely suppressed in the AM INS+GCG group during the PM HIHG clamp, whereas it was effectively suppressed in the AM INS group. Since G6Pase activity was similar between groups, the enzymatic machinery to convert G6P back to glucose was clearly sufficient for ongoing HGP in the AM INS+GCG group and therefore not rate controlling in the AM INS group. Instead, suppression of HGP in the AM INS group was likely due to the pull of G6P into the glycogenic and glycolytic pathways, likely reflecting GCK-mediated increases in G6P (55). In contrast, the sustained HGP in the AM INS+GCG group was likely due to increased glycogenolysis, which would result from reduced GCK generated G6P (56). This flux model is consistent with the observed difference in net flux between G6P and glycogen between groups, as gluconeogenic fluxes were minimal and comparable. Importantly, flux measurements provide a more accurate reflection of hepatic glucose handling than static molecular readouts (57). Together, these results suggest that antecedent glucagon reprograms hepatic glucose flux, likely in part through effects on GCK, thereby limiting substrate availability for glycogen synthesis and glycolysis.

Several points should be considered when interpreting these findings. First, tissue analyses at fixed time points may have missed transient flux regulators or dynamic enzyme activity, since steady-state protein abundance and *in vitro* enzyme assays cannot fully capture allosteric regulation or compartmentalization of metabolite pools (6). While we have previously demonstrated in dogs that hyperglucagonemia rapidly and sustainably elevates hepatic G6P under hyperglycemic conditions (34), the dynamic intracellular G6P pools during a hyperinsulinemic-hyperglycemic clamp are difficult to capture reliably with our current assay, as rapid glucose uptake and phosphorylation can alter G6P content within minutes. Nevertheless, reductions in GCK expression, NHGU, and glycogen synthesis strongly suggest altered G6P availability in the AM INS+GCG group, consistent with mechanistic predictions. Future studies investigating subcellular compartmentalization, glycogen-associated microdomains, and allosteric regulation of glycogen enzymes will be useful for a complete understanding of hepatic glucose retention in the context of the second meal phenomenon (19, 58). Clamp studies provide the advantage of tightly regulating hormone and substrate concentrations, allowing precise assessment of their effects on hepatic glucose metabolism. However, it is important to note that our experimental design, which involved prolonged glucagon elevation for 4 hours alongside concurrent hyperinsulinemia during the morning clamp, does not fully replicate the dynamic hormonal environment elicited by a typical mixed or high-protein meal, in which glucagon typically rises and falls within one to two hours depending on meal composition (59, 60). Consequently, while our findings clarify the role of sustained glucagon exposure in reprogramming hepatic glucose flux, they may not directly predict the magnitude or timing of the second-meal effect under physiological dietary conditions. Additionally, because the classical Staub-Traugott paradigm uses identical meals, it remains unclear how these mechanisms operate when sequential meals differ, as is common across breakfast, lunch, and dinner. The canine model, with its unique vascular access to the liver and the gut, offers the flexibility to study both clamp and meal conditions. In this study, clamps were used to isolate the direct effects of morning insulin and glucagon, allowing individual contributions to be evaluated before adding the complexity of mixed meals in future studies. Additionally, while hyperglucagonemia was modeled to maintain a physiologic insulin-to-glucagon molar ratio, this approach may not fully replicate the timing, magnitude, or dynamics of glucagon release during mixed or high-protein meals. Despite significant progress, many aspects of glucagon physiology remain poorly understood, particularly its interactions with other hormones and its diverse roles in hepatic and systemic metabolic pathways (61, 62). Continued research is important to unravel these complexities and guide development of effective therapies targeting glucagon signaling.

In conclusion, morning glucagon exposure induces hepatic metabolic memory, reprogramming glucose metabolism and influencing glucose homeostasis later in the day. Specifically, morning glucagon reduced PM NHGU, limited glycogen synthesis, and blunted the suppression of PM HGP during an afternoon HIHG clamp. Mechanistically, this effect likely reflects constraints on insulin-stimulated glucose phosphorylation by GCK, which may contribute to substrate-driven limitations on hepatic glycogen storage and glycolysis. Importantly, non-hepatic glucose uptake remained unchanged across groups, suggesting that these effects were largely confined to the liver and did not impact peripheral tissue glucose disposal. These findings identify glucagon as a key regulator of hepatic glucose partitioning and highlight the critical role of early meal-associated insulin-glucagon interactions in shaping the liver’s response to subsequent meals.

## 5 Acknowledgments

### 5.1 Disclosure Statement

A.D.C. has research contracts with Alnylam and Abvance, as well as grants from the NIH. A.D.C. is a consultant to Novo Nordisk, Fractyl, Sensulin, and Thetis. No other individuals have conflicts of interest relevant to this article.

### 5.2 Author Contributions

H.W., D.S.E., and A.D.C. contributed to the design of the experiments. H.W. directed all experiments and collected data. H.W., B.F., K.Y., and T.H. participated in experimental execution. B.F. and K.Y. were responsible for surgical preparation and oversight of animal care. H.W. and M.S.S. performed biochemical and tissue analyses. K.J.B., R.M.O., and D.P.C. designed and carried out the G6Pase activity assays. H.W., D.S.E., and A.D.C. interpreted the experimental results. H.W. prepared figures and drafted the manuscript, and H.W., D.S.E., G.K., and A.D.C. edited and revised it. All authors approved the final version of the manuscript.

### 5.3 Guarantor Statement

A.D.C. is the guarantor of this work and, as such, has full access to all the data in the study and takes responsibility for the integrity of the data and the accuracy of the data analysis.

### 5.4 Funding

This work was supported by National Institutes of Health (NIH) grant R01DK131082 (to A.D.C), R01DK132259 (to R. O’B) and R01GM149686 (to D.P.C). H.W. was supported by NIH grant F31DK142297 and by the Vanderbilt University Training Program in Molecular Endocrinology NIH grant T32DK007563. Hormone analysis was completed by Vanderbilt’s Hormone Assay and Analytical Services Core, supported by NIH grants DK020593 and DK059637. Vanderbilt’s Large Animal Core provided surgical expertise, supported by the NIH Grant DK020593.

### 5.5 Prior Presentation

This work was presented at the American Diabetes Association 2024 meeting. The abstract can be viewed at doi.org/10.2337/db24-1565-P.

## Notes

### Summary of Updates

The figures have been updated to reflect full-analyses, and the discussion has been expanded.

## References

1. Abraira C, Buchanan B, Hodges L. Modification of glycogen deposition by priming glucose loads: the second-meal phenomenon. Am J Clin Nutr. 1987;45(5):952–7.

2. Jovanovic A, Gerrard J, Taylor R. The second-meal phenomenon in type 2 diabetes. Diabetes Care. 2009;32(7):1199–201.

3. Bonuccelli S, Muscelli E, Gastaldelli A, Barsotti E, Astiarraga BD, Holst JJ, et al. Improved tolerance to sequential glucose loading (Staub-Traugott effect): size and mechanisms. Am J Physiol Endocrinol Metab. 2009;297(2):E532–7.

4. Wajngot A, Grill V, Efendić S, Cerasi E. The Staub-Traugott effect. Evidence for multifactorial regulation of a physiological function. Scand J Clin Lab Invest. 1982;42(4):307–13.

5. Moore MC, Smith MS, Farmer B, Coate KC, Kraft G, Shiota M, et al. Morning Hyperinsulinemia Primes the Liver for Glucose Uptake and Glycogen Storage Later in the Day. Diabetes. 2018;67(7):1237–45.

6. Waterman HL, Moore MC, Smith MS, Farmer B, Yankey K, Scott M, et al. Improved Afternoon Hepatic Glucose Disposal and Storage Requires Morning Engagement of Hepatic Insulin Receptors. Diabetes. 2024;74(3):270–81.

7. Rojas JM, Schwartz MW. Control of hepatic glucose metabolism by islet and brain. Diabetes Obes Metab. 2014;16 Suppl 1(0 1):33-40.

8. Poggiogalle E, Jamshed H, Peterson CM. Circadian regulation of glucose, lipid, and energy metabolism in humans. Metabolism. 2018;84:11–27.

9. Lin EE, Scott-Solomon E, Kuruvilla R. Peripheral Innervation in the Regulation of Glucose Homeostasis. Trends Neurosci. 2021;44(3):189–202.

10. Ramnanan CJ, Edgerton DS, Kraft G, Cherrington AD. Physiologic action of glucagon on liver glucose metabolism. Diabetes Obes Metab. 2011;13 Suppl 1(Suppl 1):118–25.

11. Jiang G, Zhang BB. Glucagon and regulation of glucose metabolism. Am J Physiol Endocrinol Metab. 2003;284(4):E671–8.

12. Hatting M, Tavares CDJ, Sharabi K, Rines AK, Puigserver P. Insulin regulation of gluconeogenesis. Ann N Y Acad Sci. 2018;1411(1):21–35.

13. Alsalim W, Tura A, Pacini G, Omar B, Bizzotto R, Mari A, et al. Mixed meal ingestion diminishes glucose excursion in comparison with glucose ingestion via several adaptive mechanisms in people with and without type 2 diabetes. Diabetes Obes Metab. 2016;18(1):24–33.

14. Holste LC, Connolly CC, Moore MC, Neal DW, Cherrington AD. Physiological changes in circulating glucagon alter hepatic glucose disposition during portal glucose delivery. Am J Physiol. 1997;273(3 Pt 1):E488–96.

15. Rivera N, Ramnanan CJ, An Z, Farmer T, Smith M, Farmer B, et al. Insulin-induced hypoglycemia increases hepatic sensitivity to glucagon in dogs. J Clin Invest. 2010;120(12):4425–35.

16. Park YM, Heden TD, Liu Y, Nyhoff LM, Thyfault JP, Leidy HJ, et al. A high-protein breakfast induces greater insulin and glucose-dependent insulinotropic peptide responses to a subsequent lunch meal in individuals with type 2 diabetes. J Nutr. 2015;145(3):452–8.

17. Lund A, Bagger JI, Christensen M, Knop FK, Vilsbøll T. Glucagon and type 2 diabetes: the return of the alpha cell. Curr Diab Rep. 2014;14(12):555.

18. Fonseca VA. Defining and characterizing the progression of type 2 diabetes. Diabetes Care. 2009;32 Suppl 2(Suppl 2):S151–6.

19. Han HS, Kang G, Kim JS, Choi BH, Koo SH. Regulation of glucose metabolism from a liver-centric perspective. Exp Mol Med. 2016;48(3):e218.

20. Hunter AL, Bechtold DA. The metabolic significance of peripheral tissue clocks. Communications Biology. 2025;8(1):497.

21. Dong H, Sun Y, Nie L, Cui A, Zhao P, Leung WK, et al. Metabolic memory: mechanisms and diseases. Signal Transduction and Targeted Therapy. 2024;9(1):38.

22. Chen Z, Natarajan R. Epigenetic modifications in metabolic memory: What are the memories, and can we erase them? Am J Physiol Cell Physiol. 2022;323(2):C570–c82.

23. Hinte LC, Castellano-Castillo D, Ghosh A, Melrose K, Gasser E, Noé F, et al. Adipose tissue retains an epigenetic memory of obesity after weight loss. Nature. 2024;636(8042):457–65.

24. Cottam MA, Caslin HL, Winn NC, Hasty AH. Multiomics reveals persistence of obesity-associated immune cell phenotypes in adipose tissue during weight loss and weight regain in mice. Nat Commun. 2022;13(1):2950.

25. Sharples AP, Turner DC. Skeletal muscle memory. Am J Physiol Cell Physiol. 2023;324(6):C1274–c94.

26. Pilotto AM, Turner DC, Mazzolari R, Crea E, Brocca L, Pellegrino MA, et al. Human skeletal muscle possesses an epigenetic memory of high-intensity interval training. Am J Physiol Cell Physiol. 2025;328(1):C258–c72.

27. Rui L. Energy metabolism in the liver. Compr Physiol. 2014;4(1):177–97.

28. Infante M, Baidal DA, Rickels MR, Fabbri A, Skyler JS, Alejandro R, et al. Dual-hormone artificial pancreas for management of type 1 diabetes: Recent progress and future directions. Artif Organs. 2021;45(9):968–86.

29. Domínguez-Oliva A, Hernández-Ávalos I, Martínez-Burnes J, Olmos-Hernández A, Verduzco-Mendoza A, Mota-Rojas D. The Importance of Animal Models in Biomedical Research: Current Insights and Applications. Animals (Basel). 2023;13(7).

30. Cherrington AD. Banting Lecture 1997. Control of glucose uptake and release by the liver in vivo. Diabetes. 1999;48(5):1198–214.

31. Myers SR, McGuinness OP, Neal DW, Cherrington AD. Intraportal glucose delivery alters the relationship between net hepatic glucose uptake and the insulin concentration. J Clin Invest. 1991;87(3):930–9.

32. Moore MC, Coate KC, Winnick JJ, An Z, Cherrington AD. Regulation of hepatic glucose uptake and storage in vivo. Adv Nutr. 2012;3(3):286–94.

33. Waterman HL, Moore MC, Smith MS, Farmer B, Scott M, Edgerton DS, et al. Duration of morning hyperinsulinemia determines hepatic glucose uptake and glycogen storage later in the day. Am J Physiol Endocrinol Metab. 2024;327(5):E655–e67.

34. Coate KC, Ramnanan CJ, Smith M, Winnick JJ, Kraft G, Irimia-Dominguez J, et al. Integration of metabolic flux with hepatic glucagon signaling and gene expression profiles in the conscious dog. Am J Physiol Endocrinol Metab. 2024;326(4):E428–e42.

35. Nelson N. A PHOTOMETRIC ADAPTATION OF THE SOMOGYI METHOD FOR THE DETERMINATION OF GLUCOSE. Journal of Biological Chemistry. 1944;153(2):375–80.

36. Edgerton DS, Cardin S, Emshwiller M, Neal D, Chandramouli V, Schumann WC, et al. Small Increases in Insulin Inhibit Hepatic Glucose Production Solely Caused by an Effect on Glycogen Metabolism. Diabetes. 2001;50(8):1872–82.

37. Keppler D, Decker K. Methods of Enzymatic Analysis, Edited by Bergmeyer, HU. Academic Press, New York and London; 1974.

38. Hawes EM, Rahim M, Haratipour Z, Orun AR, O’Rourke ML, Oeser JK, et al. Biochemical and metabolic characterization of a G6PC2 inhibitor. Biochimie. 2024;222:109–22.

39. Bergman RN, Phillips LS, Cobelli C. Physiologic evaluation of factors controlling glucose tolerance in man: measurement of insulin sensitivity and beta-cell glucose sensitivity from the response to intravenous glucose. J Clin Invest. 1981;68(6):1456–67.

40. Shah AM, Wondisford FE. Tracking the carbons supplying gluconeogenesis. J Biol Chem. 2020;295(42):14419–29.

41. Redant S, Hussein H, Mugisha A, Attou R, De Bels D, Honore PM, et al. Differentiating Hyperlactatemia Type A From Type B: How Does the Lactate/pyruvate Ratio Help? J Transl Int Med. 2019;7(2):43–5.

42. Moore MC, Satake S, Lautz M, Soleimanpour SA, Neal DW, Smith M, et al. Nonesterified fatty acids and hepatic glucose metabolism in the conscious dog. Diabetes. 2004;53(1):32–40.

43. Satake S, Moore MC, Igawa K, Converse M, Farmer B, Neal DW, et al. Direct and indirect effects of insulin on glucose uptake and storage by the liver. Diabetes. 2002;51(6):1663–71.

44. Maki KC, Phillips-Eakley AK, Smith KN. The Effects of Breakfast Consumption and Composition on Metabolic Wellness with a Focus on Carbohydrate Metabolism. Adv Nutr. 2016;7(3):613s–21s.

45. Petersen MC, Vatner DF, Shulman GI. Regulation of hepatic glucose metabolism in health and disease. Nat Rev Endocrinol. 2017;13(10):572–87.

46. Ma X, Chen Q, Pu Y, Guo M, Jiang Z, Huang W, et al. Skipping breakfast is associated with overweight and obesity: A systematic review and meta-analysis. Obes Res Clin Pract. 2020;14(1):1–8.

47. Wikarek T, Kocełak P, Owczarek AJ, Chudek J, Olszanecka-Glinianowicz M. Effect of dietary macronutrients on postprandial glucagon and insulin release in obese and normal-weight women. Int J Endocrinol. 2020;2020:4603682.

48. Kalra S, Gupta Y. The Insulin:Glucagon Ratio and the Choice of Glucose-Lowering Drugs. Diabetes Ther. 2016;7(1):1–9.

49. Iynedjian PB, Jotterand D, Nouspikel T, Asfari M, Pilot PR. Transcriptional induction of glucokinase gene by insulin in cultured liver cells and its repression by the glucagon-cAMP system. J Biol Chem. 1989;264(36):21824–9.

50. Ramnanan CJ, Edgerton DS, Rivera N, Irimia-Dominguez J, Farmer B, Neal DW, et al. Molecular characterization of insulin-mediated suppression of hepatic glucose production in vivo. Diabetes. 2010;59(6):1302–11.

51. Agius L. Hormonal and Metabolite Regulation of Hepatic Glucokinase. Annu Rev Nutr. 2016;36:389–415.

52. Cullen KS, Al-Oanzi ZH, O’Harte FP, Agius L, Arden C. Glucagon induces translocation of glucokinase from the cytoplasm to the nucleus of hepatocytes by transfer between 6-phosphofructo 2-kinase/fructose 2,6-bisphosphatase-2 and the glucokinase regulatory protein. Biochim Biophys Acta. 2014;1843(6):1123–34.

53. Agius L. Glucokinase and molecular aspects of liver glycogen metabolism. Biochemical Journal. 2008;414(1):1–18.

54. Velho G, Petersen KF, Perseghin G, Hwang JH, Rothman DL, Pueyo ME, et al. Impaired hepatic glycogen synthesis in glucokinase-deficient (MODY-2) subjects. J Clin Invest. 1996;98(8):1755–61.

55. Abu Aqel Y, Alnesf A, Aigha, II, Islam Z, Kolatkar PR, Teo A, et al. Glucokinase (GCK) in diabetes: from molecular mechanisms to disease pathogenesis. Cell Mol Biol Lett. 2024;29(1):120.

56. Aiston S, Andersen B, Agius L. Glucose 6-phosphate regulates hepatic glycogenolysis through inactivation of phosphorylase. Diabetes. 2003;52(6):1333–9.

57. Bar-Peled L, Kory N. Principles and functions of metabolic compartmentalization. Nat Metab. 2022;4(10):1232–44.

58. Habashy A, Acree C, Kim K-Y, Zahraei A, Dufresne M, Phan S, et al. Spatial patterns of hepatocyte glucose flux revealed by stable isotope tracing and multi-scale microscopy. Nature Communications. 2025;16(1):5850.

59. Ichikawa R, Takano K, Fujimoto K, Kobayashi M, Kitamura T, Shichiri M, et al. Robust increase in glucagon secretion after oral protein intake, but not after glucose or lipid intake in Japanese people without diabetes. J Diabetes Investig. 2023;14(10):1172–4.

60. Kabadi UM. Dose-kinetics of pancreatic &#x3b1;- and &#x3b2;-cell responses to a protein meal in normal subjects. Metabolism - Clinical and Experimental. 1991;40(3):236–40.

61. Capozzi ME, D’Alessio DA, Campbell JE. The past, present, and future physiology and pharmacology of glucagon. Cell Metab. 2022;34(11):1654–74.

62. Finan B, Capozzi ME, Campbell JE. Repositioning Glucagon Action in the Physiology and Pharmacology of Diabetes. Diabetes. 2020;69(4):532–41.

