## Supplementary Materials for "Morning Glucagon Disrupts Insulin Induced Hepatic Metabolic Memory and Subsequent Afternoon Glucose Metabolism in Canines"

**Hepatic glycogen content**

Liver glycogen content was measured using a modification of the method of Keppler and Decker (1, 2). Frozen liver tissue (~180 mg) was weighed under liquid nitrogen to prevent thawing and enzymatic degradation. Tissue weight (mg) was used to calculate the volume of 0.6 N perchloric acid (PCA) required for homogenization (volume = tissue weight × 5). Samples were homogenized on ice in the calculated volume of PCA. From each homogenate, 200 µL aliquots were transferred into separate tubes and neutralized with 100 µL of potassium bicarbonate. To quantify glycogen, 500 µL of amyloglucosidase solution (2 mg/mL in 0.4 M sodium acetate buffer) was added to experimental aliquots and incubated at 40°C in a shaking water bath for 2 hours to hydrolyze glycogen to glucose. Duplicate control samples lacking amyloglucosidase were processed in parallel to account for free glucose in the homogenate. Glycogen content was calculated as glucose released in enzyme-treated samples minus glucose in paired control samples.

Standards were prepared using oyster glycogen (4, 5.5, and 7 mg) dissolved in 1 mL of 0.6 N PCA and processed alongside samples. After incubation, samples were cooled on ice and centrifuged at 3,000 rpm for 5 minutes. For samples containing tracer, 500 µL of supernatant was transferred to scintillation vials for [³H]-glycogen counting using the same evaporation, reconstitution, and scintillation methods applied to plasma 3-[³H]-glucose. Remaining aliquots were analyzed for glucose concentration using an Analox GM9 glucose analyzer. All solutions (PCA, potassium bicarbonate, amyloglucosidase) were prepared fresh on the day of assay.

**RNA extraction, cDNA synthesis, and quantitative real-time PCR**

Total RNA was isolated from canine liver tissue to assess hepatic gene expression. Approximately 50 mg of frozen liver tissue was homogenized in 1 mL of Tri-reagent (Sigma-Aldrich, St. Louis, MO) according to the manufacturer’s instructions. RNA purification was performed using the Direct-zol RNA Miniprep Kit (Zymo Research, Irvine, CA) with final elution in 35 µL of nuclease-free TE buffer. RNA yield and purity were determined by spectrophotometry, with A260/A280 ratios >1.8 considered acceptable. All procedures were conducted using RNase/DNase-free reagents and consumables. First-strand cDNA was synthesized from 1 µg of total RNA using the High-Capacity cDNA Reverse Transcription Kit (Applied Biosystems, Foster City, CA) following the manufacturer’s instructions. cDNA was stored at -80°C until use. Primers for target and reference genes were designed with Beacon Designer software (Premier Biosoft, Palo Alto, CA), verified for specificity with BLAST, and further confirmed by melt curve analysis. Primer efficiency was within the range of 91%-96%.

Quantitative real-time PCR was carried out on a CFX96 Real-Time PCR Detection System (Bio-Rad, Hercules, CA) using SsoAdvanced Universal SYBR Green Supermix (Bio-Rad). Each 25 µL reaction contained 100 ng cDNA template, 12.5 µL SYBR Green Supermix, 0.4 µM forward and reverse primers, and nuclease-free water. Thermal cycling conditions were: 95°C for 3 minutes, followed by 39 cycles of 95°C for 10 seconds and 55°C for 30 seconds. All reactions were run in duplicate. Melt curve analysis was performed to ensure amplification specificity. Relative gene expression was calculated using the 2^-ΔΔCt method, with GAPDH as the reference gene (3). Data represent the mean of 2-3 independent PCR runs. Tissue from the left central and left lateral lobes was analyzed, as these lobes account for ~50% of total liver mass and provide a representative assessment of hepatic gene expression. Primer details can be viewed in Table 1 below.

**Supplemental Table 1 - Primer details for quantitative PCR of selected genes**

| **Gene Name** | **Forward Primer (5’ 🡪 3’)** | **Forward Melting Temperature (Tm)** | **Reverse Primer (5’ 🡪 3’)** | **Reverse Melting Temperature (Tm)** |
| --- | --- | --- | --- | --- |
| *GCK* | CAGAGGGGACTTTGAAATG | 59.6°C | CTGCATCTCCTCCATGTAG | 58.3°C |
| *PYGL* | TCGCAGACTATGAAGCCTATG | 61.9°C | CCTTAATTGTTOGGTCACTAGAG | 60.5°C |
| *FOXO1* | CTACGAGTGGATGGTCAAGAG | 61.3°C | CACGAATGAACTTGOTATGTAGG | 62.3°C |
| *PCK1* | AGCTTTCAATGCCCGATTTCCAGG | 73.1°C | TCAGCTCGATGCCGATCTTTGACA | 73.7°C |
| *G6PC1* | CCTTTATTCCTCTTTC | 50.7°C | GGTGTTGCTATAGTAGTC | 46.0°C |
| *SLC37A4* | GGAATTCCGGAGTCCAACATCAGCAGGTTC | 78.0°C | CCCAAGCTTGCCATCTCAGTTTGGCACTTGGTGG | 83.8°C |
| *SREBP-1c* | GTGAAGGCAGCGGGTATCAG | 66.8°C | TCTCAGTGTCCACTACCAGAGG | 63.2°C |
| *PGC1-α* | GCTTTCTGGGTGGACTCAAGTG | 67.0°C | GCAAGTTCGCCTCGTTCTCTTC | 68.2°C |
| *FAS* | TACTGGAGGGGCCAGTGCATCA | 72.9°C | GTCCCGAGATGGTCACTGTGTC | 68.1°C |
| *GAPDH* | TGTCCCCACCCCCAATGTATC | 69.8°C | CTCCGATGCCTGCTTCACTACCTT | 70.2°C |

**Western blotting**

Frozen liver tissue (~100 mg) was homogenized in 1 mL of ice-cold homogenization buffer (20 mM Tris, 200 mM NaCl, 50 mM NaF, 1 mM EDTA, 1 mM EGTA, 10% glycerol, 1% SDS, pH 7.2) supplemented with protease and phosphatase inhibitors (Sigma-Aldrich, St. Louis, MO). Homogenates were centrifuged at 3,000 × g for 10 minutes at 4°C, and supernatants were collected. Protein concentrations were determined using a BCA protein assay (Bio-Rad, Hercules, CA). Protein samples were adjusted to 4 µg/µL in Laemmli buffer, heat-denatured at 95°C for 5 minutes, and stored at -80°C. Equal amounts of protein (40 µg per lane) were separated on 4-12% Criterion TGX gels (Bio-Rad) and transferred to nitrocellulose membranes using a semi-dry Trans-Blot SD transfer system (Bio-Rad) in Towbin buffer (25 mM Tris, 192 mM glycine, 20% methanol, pH 8.3) for 25 minutes at 15 V. Membranes were stained with Ponceau S solution to visualize total protein, imaged with a ChemiDoc system (Bio-Rad), and normalized to Ponceau S staining intensity. Membranes were then washed with TBST (10 mM Tris, 150 mM NaCl, 0.2% Tween-20, pH 7.5) and blocked for 1 hour at room temperature with 5% (w/v) bovine serum albumin in TBST. Blots were incubated overnight at 4°C with primary antibodies diluted in blocking buffer. The following primary antibodies were used in western blotting analysis: pAkt (#9271, Cell Signaling Technology, Danvers, MA), tAkt (#9272, Cell Signaling Technology), pGS (#98348, Cell Signaling Technology), tGS (#3893, Cell Signaling Technology), pGP (#ab227043, Abcam, Cambridge, UK), tGP (#ab198268, Abcam), pFOXO1 (#9461, Cell Signaling Technology), tFOXO1 (#2880, Cell Signaling Technology), and GK (#sc-17819, Santa Cruz Biotechnology). Primary antibody dilutions were all 1:5,000, except for GK (1:10,000). pFOXO1 (1:1,000), and tFOXO1 (1:1,000). After three washes in TBST (x5 min), membranes were incubated with 1:5,000 Anti-rabbit HRP-conjugated secondary antibodies (Promega, Madison, WI) for 1 hour at room temperature, followed by three additional washes (x5 min) in TBST. Proteins were detected using ECL Plus reagents (GE Healthcare, Piscataway, NJ) and imaged with a ChemiDoc system. Band intensities were quantified with ImageJ software (NIH, Bethesda, MD), normalized to total protein detected by Ponceau S. Data represent the mean of two blots per protein. Western blotting was performed on tissue from the left central and left lateral lobes, which together account for ~50% of total liver mass in the dog.

**Glucose-6-phosphatase (G6Pase) activity assay**

**Isolation of microsomal membranes**

Microsomal membranes were isolated from frozen dog liver as previously described (4). Briefly, ~1.0 g of frozen liver was added to 10 mL of a 50 mM Hepes pH 7.4, 25 mM KCl, 5 mM MgCl2, 0.25 M sucrose solution and homogenized with three strokes of a motorized dounce. Non-homogenized tissue was removed by centrifuging the homogenate at 7800 g for 6 min at 4°C. The supernatant was then spun at 214,000 g for 30 min at 4 C using a Beckman TLA 100.3 rotor to isolate a microsomal membrane fraction. This fraction was resuspended in 900 mL of a 29 mM MES, 21 mM Tris, 50 mM NaCl pH 6.5 solution.

**Measurement of G6Pase activity in vitro**

G6Pase activity present in microsomal membranes isolated from liver was quantified as previously described (4). Total microsomal protein was measured using a BioRad Bradford Assay. 40 mg of microsomal protein was mixed with reaction buffer (58 mM MES, 42 mM Tris, 100 mM NaCl pH 6.5) supplemented with either 10 mM G6P or 10 mM M6P. Reactions were incubated in a 30°C water bath for 15 min and then placed on ice for 1 min. The reactions were quenched using a 12% SDS solution. A cold inorganic phosphate (P_i_) chelating solution (1:1 2% ammonium molybdate:12% ascorbic acid solution) was added and reactions incubated for 5 min at room temperature, followed by addition of a developing solution (80 mM sodium citrate, 150 mM sodium meta-arsenite, 2% glacial acetic acid) and a 20 min incubation at room temperature. The absorbance of the reactions was measured using a plate reader at 850 nm. A Pi standard curve was used to calculate the total nmol Pi released based on the absorbance at 850nm. Background Pi from the buffers and microsomes in the absence of G6P or M6P was subtracted to calculate total Pi generated. G6Pase activity was determined by dividing the total Pi generated by the length of the assay and the total microsomal protein present.

**Glycogen synthase enzymatic activity assay**

Glycogen synthase (GS) activity was assayed by incorporation of [¹⁴C]-UDP-glucose into glycogen, adapted from Shiota (2). Stock solutions of Tris-HCl (400 mM, pH 7.0 or 8.5) and EDTA (100 mM, pH 7.0 or 8.5) were prepared in advance and stored at room temperature. UDP-[^14^C]-glucose was aliquoted and dried at –20°C, then reconstituted with unlabeled UDP-glucose on the day of assay. Glucose-6-phosphate (G6P) solutions were prepared fresh. Homogenization buffer (100 mM potassium fluoride, 10 mM EDTA, 1% glycogen, pH 7.0) was used to prepare liver extracts. Dilution buffers containing potassium fluoride, EDTA, and glycogen were adjusted to either pH 7.0 (Buffer A) or pH 8.5 (Buffer B) to assess GS activity in different phosphorylation states. Reaction mixtures contained 50 mM Tris (pH 7.0 or 8.5), 15 mM EDTA, 1% glycogen, 18 mM UDP-[^14^C]-glucose, and either low (0.375 mM) or high (18 mM) G6P. Low G6P reflects “active” synthase activity, whereas high G6P allows determination of both active and total activity.

Approximately 100 mg of frozen liver tissue was homogenized in 1 mL of homogenization buffer on ice. Homogenates were centrifuged at 500 × g for 10 minutes at 4°C, and supernatants were diluted tenfold in pre-warmed dilution buffer A or B. Diluted homogenates were equilibrated at 37°C for 10 minutes prior to assay. Reactions were initiated by adding 100 µL of homogenate to 200 µL of pre-warmed reaction buffer. At 0, 10, and 20 minutes, 50 µL aliquots were spotted onto filter paper (20 × 25 mm), dried briefly, and transferred to 70% ethanol with stirring to precipitate glycogen and remove unincorporated substrate. Filters were washed sequentially with 70% ethanol (3 × 10 minutes), followed by 100% ethanol (5 minutes), and air-dried overnight. Dried filters were placed in scintillation vials with 1 mL water and 10 mL scintillation fluid, mixed, stored for 24 hours, and analyzed by liquid scintillation counting. Parallel control reactions lacking tissue homogenate were performed to measure background UDP-[^14^C]-glucose incorporation.

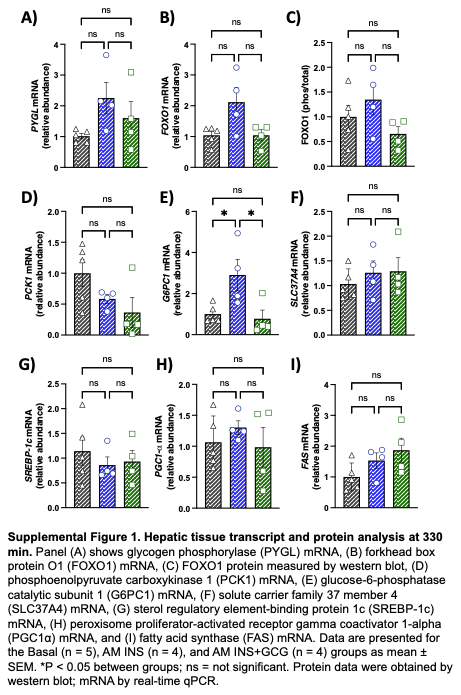

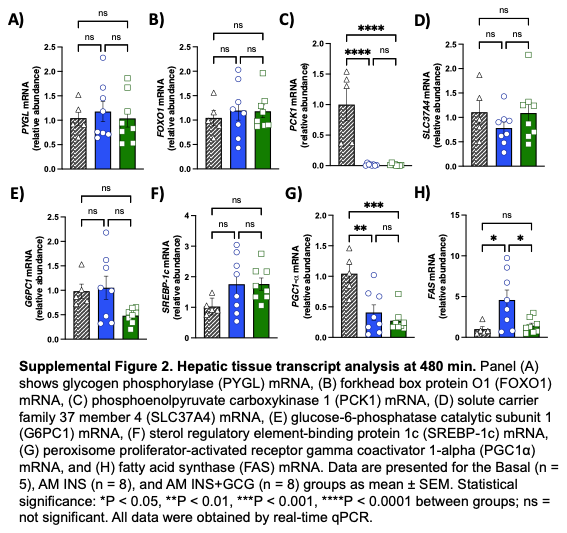

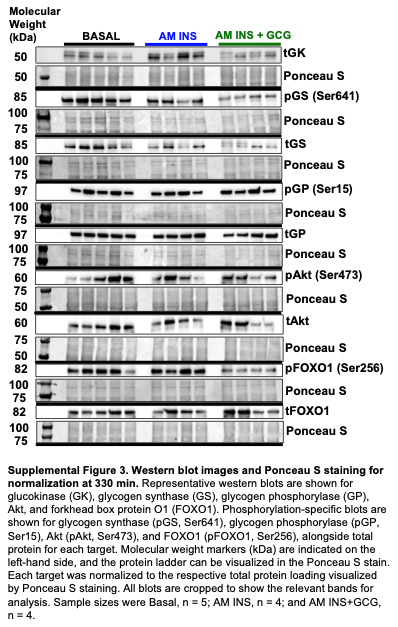

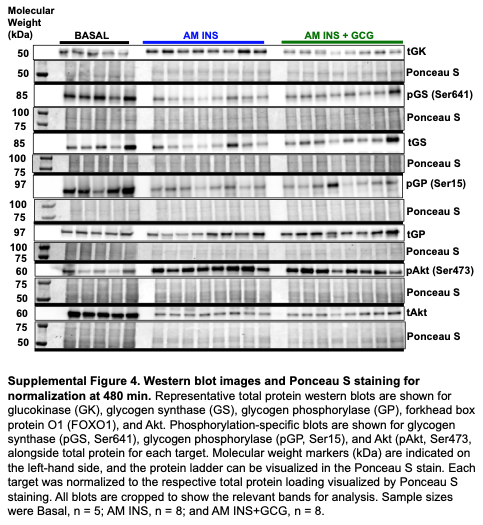

References

1.	Keppler D DK. Glycogen determination with amyloglucosidase. *Methods of Enzymatic Analysis.* 1974;3:1127-31.

2.	Keppler D, and Decker K. Academic Press, New York and London; 1974.

3.	Livak KJ, and Schmittgen TD. Analysis of relative gene expression data using real-time quantitative PCR and the 2(-Delta Delta C(T)) Method. *Methods.* 2001;25(4):402-8.

4.	Hawes EM, Rahim M, Haratipour Z, Orun AR, O'Rourke ML, Oeser JK, et al. Biochemical and metabolic characterization of a G6PC2 inhibitor. *Biochimie.* 2024;222:109-22.
